# A conserved transcriptional axis links cortical organization, developmental timing and autism-associated perturbations

**DOI:** 10.64898/2026.09.05.749572

**Authors:** Shangzheng Huang, Yingjie Peng, Xiaohan Tian, Changsheng Dong, Tongyu Zhang, Yingchao Shi, Ang Li

## Abstract

Much of our mechanistic understanding of cortical development and neurodevelopmental disorders comes from studies in mice, yet translating these insights to humans rests on a fundamental assumption: that the cortex is organized according to conserved principles across species. Here, by independently decomposing human and mouse cortical transcriptomes, we test this assumption and identify a shared organizational axis extending from limbic anterior–ventral (AV) to primary sensory posterior–dorsal (PD) cortex. This axis aligns with conserved variation in cell-type composition, thalamocortical connectivity, myelination and excitation–inhibition balance. Its spatial topology emerges by mid-gestation and is progressively refined while remaining stable in orientation across subsequent development. What varies systematically along the axis is developmental timing: AV-enriched genes preferentially retained earlier cortical-construction features and progressively decline after birth, whereas PD-enriched genes preferentially reflected later maturation processes and progressively increased, with developmental rates graded along the axis. Across genetically distinct autism mouse models, developmental dysregulation converged on this axis, following a shared pattern that generally intensified toward the PD pole. The same axis also organized genotype-specific patterns of cortical volume alteration, while human autism risk genes showed corresponding enrichment along this transcriptional coordinate. Overall, we identify a conserved transcriptional axis that links cortical architecture to developmental timing and organizes the heterogeneous cortical effects of autism-associated mutations.

## Main

The mammalian cortex is built through the spatially patterned and temporally coordinated expression of more than 20,000 genes^1–3^. Comparative studies have identified many conserved elements of cortical development^4^, including core transcription factors^5,6^, Wnt^7^ and Notch^8,9^ signaling pathways, and broadly shared cortical cell types^10–12^. Yet whether these conserved elements are assembled into a common, cortex-wide organizational program remains unknown. Spatial transcriptional gradients provide a tractable way to test this possibility: by capturing continuous molecular variation across the cortical sheet, they allow us to ask whether homologous genes, cell types and developmental processes are aligned along a common coordinate across species.

In the adult human cortex, a major axis of macroscale organization extends from primary sensorimotor to transmodal association regions^13–16^. This sensorimotor–association (S–A) axis aligns with multiple brain phenotypes, including the principal gradient of functional connectivity^17^, patterns of human cortical expansion relative to macaques^18^, and the cortical distribution of the T1-weighted to T2-weighted (T1w/T2w) MRI signal ratio, a putative marker of cortical myelination^15^. Comparable gradients have also been recovered de novo in macaque and marmoset cortex^15,19^. Extending this framework to rodents, however, is not straightforward because some of the association areas anchoring its transmodal pole in primates—particularly granular lateral prefrontal and inferior parietal cortex—lack clear homologues in rodents^20,21^. The S–A axis may therefore capture an important dimension of primate cortical specialization without necessarily providing the most appropriate coordinate for testing conserved mammalian organization.

Recent transcriptomic decomposition studies have further dissociated limbic-to-sensory and frontal– occipital–temporal gradients from the canonical S-A axis in the human cortex^22–24^. A corresponding organization has also been inferred in mouse cortex by projecting human-derived gene weights onto mouse expression data^24^. Together, these observations raise the possibility of a conserved organizational coordinate of the mammalian cortex. However, projection of a human-derived axis into mouse data cannot establish that the same gradient can be recovered independently in both species. More importantly, similar spatial topographies do not necessarily imply a shared underlying program: each lineage could, in principle, assemble a similar-looking gradient from different genes, cell types and molecular processes.

Moreover, establishing a conserved cortical organization program requires more than demonstrating cross-species spatial correspondence. If one axis is the spatial readout of a conserved gene-defined developmental program, its topology should emerge during cortical development, and position along the axis should be systematically related to the temporal trajectories of the genes that construct it. Whether these spatial and temporal features are conserved between humans and mice remains unknown. This question is consequential because much of the mechanistic understanding of cortical development and its disorders derives from mouse models and is extrapolated to humans, implicitly assuming that the relevant cortical programs are conserved across species. Autism spectrum disorder (ASD) provides a particularly stringent test of this framework: despite substantial genetic heterogeneity, distinct ASD-associated mutations may disrupt common developmental processes. Whether their molecular and anatomical effects converge along a conserved cortical developmental coordinate is also not well characterized.

Here we integrated whole-cortex spatial transcriptomic datasets from mouse cortex with human cortical transcriptomic resources spanning mid-gestation to adulthood^13,25–30^. We derived the major transcriptional axes independently within each species and registered them using TransBrain^24^, a quantitative mouse– human brain translation framework. Among the gradients examined, only one showed significant cross-species correspondence, extending from limbic anterior–ventral cortex to primary sensory posterior–dorsal cortex; we therefore refer to it as the AV–PD axis. Developmentally, the spatial orientation of this axis was evident by mid-gestation and was progressively refined without reorientation in both species (**Fig. 1a**). Along this stable spatial scaffold, genes toward its opposing poles followed distinct, continuously graded postnatal trajectories. Specifically, AV-pole genes were preferentially expressed prenatally and declined after birth, retaining more cortical construction features, whereas PD-pole genes progressively increased toward adulthood, reflecting later maturation processes, with developmental rates graded continuously along the axis (**Fig.1b**). The proportions of homologous cortical cell types showed corresponding distributions along the axis across primate spatial transcriptomic data^31^, multi-donor human single-nucleus RNA-sequencing data^32^, whole-brain mouse spatial atlases^29,30^ and complementary cell-marker^33^ enrichment analyses (**Fig. 1c**). The axis also aligned across species with thalamocortical projection gradients^34^, T1w/T2w maps^15,35^ and excitation–inhibition balance^36^ (**Fig. 1d**). Finally, developmental transcriptional dysregulation across genetically distinct ASD mouse models^37^ followed a shared AV–PD axis-dependent pattern that intensified toward the PD pole. Cortical volume alterations across ASD-related mouse genotypes^38^ were also organized along this coordinate, while autism-associated signals from genome-wide association^39,40^, differential-expression^41–46^ and rare risk genes (from SFARI^47^) also showed corresponding non-random distributions along the axis (**Fig. 1e)**. Together, these findings establish the AV– PD axis as a conserved spatial readout of a gene-defined developmental program that links cortical architecture to developmental timing and organizes the heterogeneous cortical effects of autism-associated mutations.

**Figure 1.**
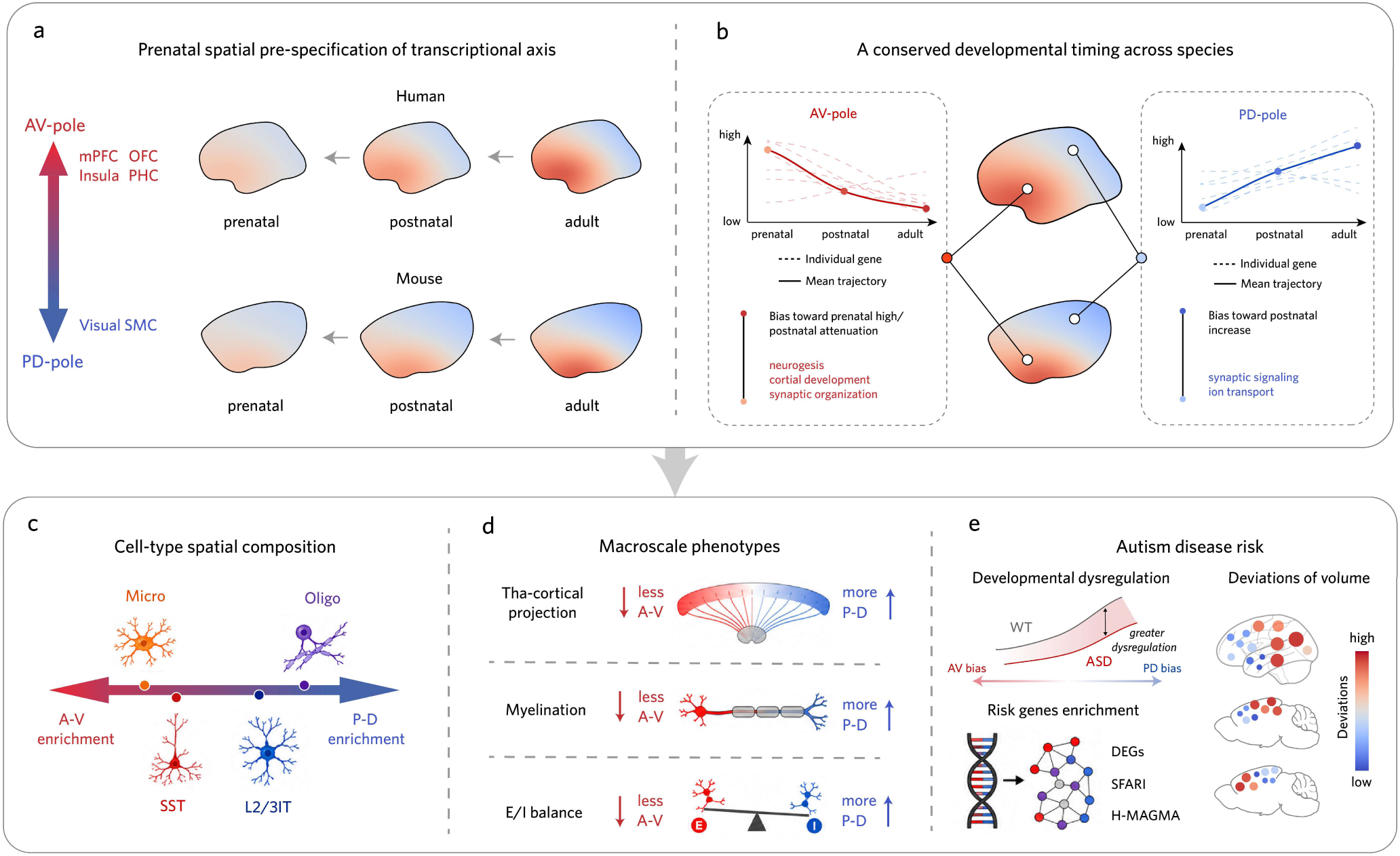
Summary of the cross-species AV–PD axis findings. **(a)** The AV–PD axis extends from anterior–ventral limbic cortices (AV pole; red: mPFC, OFC, insula and PHC) to posterior–dorsal sensory cortices (PD pole; blue: visual, and sensorimotor cortices). An adult-like spatial configuration is already evident prenatally and becomes progressively accentuated during postnatal maturation in both humans (top) and mice (bottom). **(b)** Conserved pole-specific developmental trajectories across species. AV-pole genes (red) show high prenatal expression with postnatal downregulation, and are enriched for neurogenesis, cortical development, and synaptic organization; PD-pole genes (blue) show the opposite trajectory and are enriched for synaptic signaling and ion transport. **(c)** Cell-type composition varies systematically along the axis, with microglia and SST interneurons enriched toward the AV pole, oligodendrocytes and L2/3 IT neurons enriched toward the PD pole. **(d)** Cross-species macroscale phenotypes—including thalamocortical projection strength, myelination and the E/I ratio—show concordant gradients, with higher values toward the PD pole. **(e)** Autism-related molecular, anatomical, and genetic signals converge along the AV–PD axis. AV, anterior–ventral; PD, posterior–dorsal; mPFC, medial prefrontal cortex; OFC, orbitofrontal cortex; PHC, parahippocampal cortex; SMC, sensorimotor cortex; SST, somatostatin-expressing interneurons; L2/3 IT, layer 2/3 intratelencephalic; E/I, excitation-to-inhibition; DEGs, differentially expressed genes; SFARI, Simons Foundation Autism Research Initiative; H-MAGMA, Hi-C-coupled MAGMA.

## Results

### Independent analyses identify a conserved AV–PD transcriptional axis

Previous work using region-specific transcriptional embedding identified an AV–PD axis as a major transcriptional component of the human cortex (variance explained = 26.8%), and a spatially comparable pattern was inferred in mouse cortex by projecting human-derived gene weights onto mouse data^24^ (**Fig. 2a**). However, whether this axis represents an intrinsic component of mouse cortical organization or a pattern imposed by cross-species projection remains unknown.

**Figure 2.**
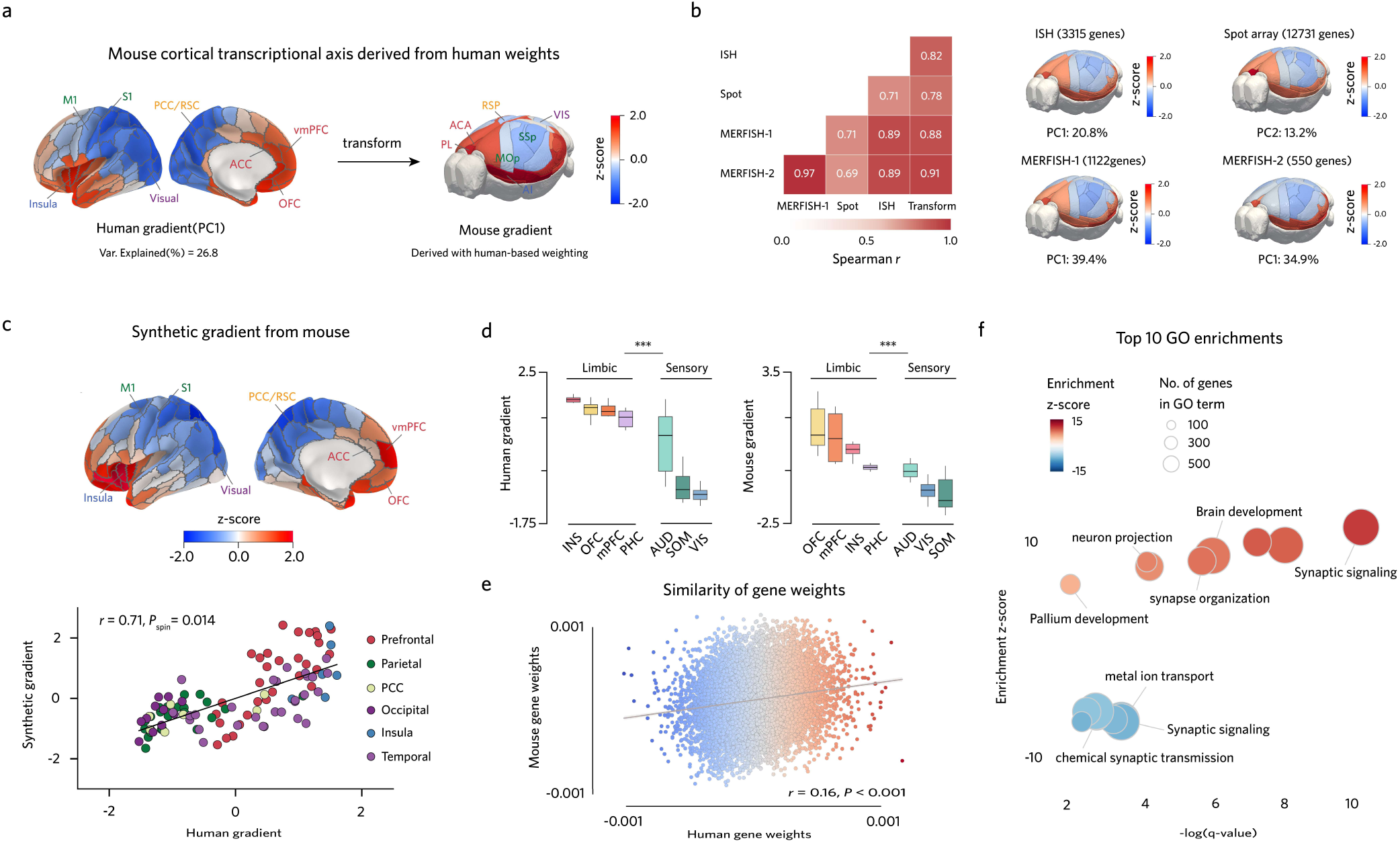
Independent recovery of the AV–PD transcriptional axis in humans and mice. **(a)** Principal cortical transcriptional gradient (PC1) in the adult human cortex (left; variance explained = 26.8%) and its projection onto the mouse^24^ (right), both along the AV–PD axis. Colors indicate z-scores. **(b)** PCA decomposition of mouse transcriptomic datasets. Left, correlations among four independently derived transcriptional components (ISH^27^; Spot array^28^; MERFISH-1^29^; MERFISH-2^30^) and the human-derived projection^24^ (Transform; Spearman *r* = 0.69–0.97, all FDR-adjusted *P* < 0.001, *n* = 39 ROIs). Right, visualization of the component in each dataset (variance explained: ISH PC1, 20.8%; Spot PC2, 13.2%; MERFISH-1 PC1, 39.4%; MERFISH-2 PC1, 34.9%). **(c)** Cross-species correspondence between the TransBrain^24^-derived synthetic human map (top) and the human gradient (bottom; Spearman *r* = 0.71, *P_spin_* = 0.014, *n* = 105 ROIs, 1,000 spatial rotations). Points are colored by lobe. **(d)** AV–PD transition from limbic to sensory cortices in humans and mice (human: rank-biserial *r* = 0.85, *P* = 1.47 × 10^−9^*, n* = 25 limbic and 56 sensory ROIs; mouse: *r* = 0.92, *P* = 5.93 × 10^−6^*, n* = 14 and 21 ROIs, respectively; two-sided Mann–Whitney *U* test). Box plots: median, 25th–75th percentiles; whiskers, 1.5 × IQR. **(e)** Cross-species correspondence of independently derived gene weights between humans (AHBA) and mice (Spot array; Pearson *r* = 0.16, *P* = 7.15 × 10^−^⁷³, *n* = 12,731 genes). Points are colored by human gene weights (red, AV-biased; blue, PD-biased). Shaded band, 95% confidence interval of linear fit. **(f)** Top 10 enriched GO biological-process terms of significant cross-species AV–PD-related genes. The x-axis shows enrichment significance (-log(q), FDR-corrected), and the y-axis enrichment z-score (positive: 442 AV-pole genes; negative: 428 PD-pole genes). Bubble size, number of genes per term. Brain region abbreviations are defined in **Supplementary Table 3.** \*\*\**P* < 0.001.

Therefore, we analyzed four independent whole-cortex spatial transcriptomic datasets: an ISH dataset^27^, a spot-array dataset^28^, and two MERFISH datasets (MERFISH 1^29^ and MERFISH 2^30^). PCA decomposition was applied to each dataset (**Methods**) to identify major cortical transcriptional gradients. Across datasets, one transcriptional component was consistently recovered, corresponding to PC1 in three datasets and PC2 in the spot-array dataset (13.2–39.4% variance explained), with high spatial concordance (mean Spearman *r* = 0.81; **Fig. 2b**; **Extended Data Fig. 1a**).

We next asked whether this mouse-intrinsic component corresponds to the AV–PD axis identified in human cortex. Three complementary analyses supported cross-species conservation. First, the independently recovered mouse component matched the previously inferred mouse pattern obtained by projecting human-derived gene weights^24^ (mean Spearman *r* = 0.85; **Fig. 2b**). Second, after cross-species mapping using TransBrain^24^, the mouse gradient showed significant correspondence with the independently derived human cortical gradient (Spearman *r* = 0.71, *P_spin_* = 0.014, n = 1,000 rotations; **Fig. 2c**), with consistent limbic-to-sensory transitions across species (human: rank-biserial *r* = 0.85, *P* = 1.47×10^−9^; mouse: *r* = 0.92, *P* = 5.93×10^−6^; **Fig. 2d**; **Extended Data Fig. 1b, c**). Third, homologous gene contributions to the gradient were significantly correlated between species (Pearson *r* = 0.16, *P* = 7.15×10^−^⁷³; **Fig. 2e**), supporting partial conservation of the molecular determinants underlying the AV–PD coordinate.

We next identified AV–PD axis genes that were significantly associated with the gradient in both humans and mice. Gene Ontology (GO) biological process enrichment revealed distinct programs at the two poles (**Methods**). AV-pole genes were predominantly enriched for developmental and structural programs, including brain development, neuron projection, axon development, synaptic organization and synaptic signaling. In contrast, PD-pole genes were predominantly enriched for activity-related programs, including chemical synaptic transmission, metal ion transport and synaptic signaling (**Fig. 2f**; **Supplementary Fig. 1**). Although both poles converged on synaptic categories, the underlying sub-programs were distinct: AV-pole genes were preferentially enriched for synaptic organization, whereas PD-pole genes were preferentially enriched for synaptic transport (**Supplementary Fig. 2**).

Because multiple transcriptional gradients coexist in the human cortex, we also compared the first three mouse transcriptional components with three major human transcriptional gradients previously identified^22–24^. Only the AV–PD axis showed significant cross-species correspondence (**Extended Data Fig. 1d**). We also examined whether the AV–PD transcriptional axis reflected a widely reported canonical sensorimotor S–A hierarchy, a low-dimensional organization derived from functional connectivity^17^. The human AV–PD gradient was not significantly aligned with the functional S–A gradient (Spearman *r* = - 0.38, *P_spin_* = 0.08; **Extended Data Fig. 1e**), suggesting that the AV–PD axis captures a molecular organization distinct from the established functional hierarchy.

Together, these results establish the AV–PD axis as an independently recovered and cross-species conserved mode of cortical transcriptional organization across mammals.

### The AV–PD axis captures conserved cellular and macroscale organization

Cortical organization emerges across multiple biological scales, from transcriptional programs and cell-type composition to circuit architecture and macroscale phenotypes^16,48,49^. Having established that the AV– PD axis represents a conserved transcriptional coordinate underlying cortical organization, we asked whether this molecular organization is reflected across cellular and macroscale levels of cortical architecture.

Given the lack of whole-cortex, spatially resolved cell-type maps in the human brain, we used a human map inferred via connectome-based embedding^50^ from macaque spatial transcriptomics^31^ and whole-brain MERFISH map in mice^29,30^ (**Methods**). Across species, cortical cell-type composition distributions showed conserved polarity along the AV–PD axis (**Fig. 3a**). AV-pole regions were enriched for oligodendrocyte progenitor cells (OPCs), Sst interneurons, chandelier cells and microglia-perivascular macrophages (Micro-PVM), whereas PD-pole regions were enriched for layer 2/3 intratelencephalic projection neurons (L2/3 IT) and oligodendrocytes (Oligo).

**Figure 3.**
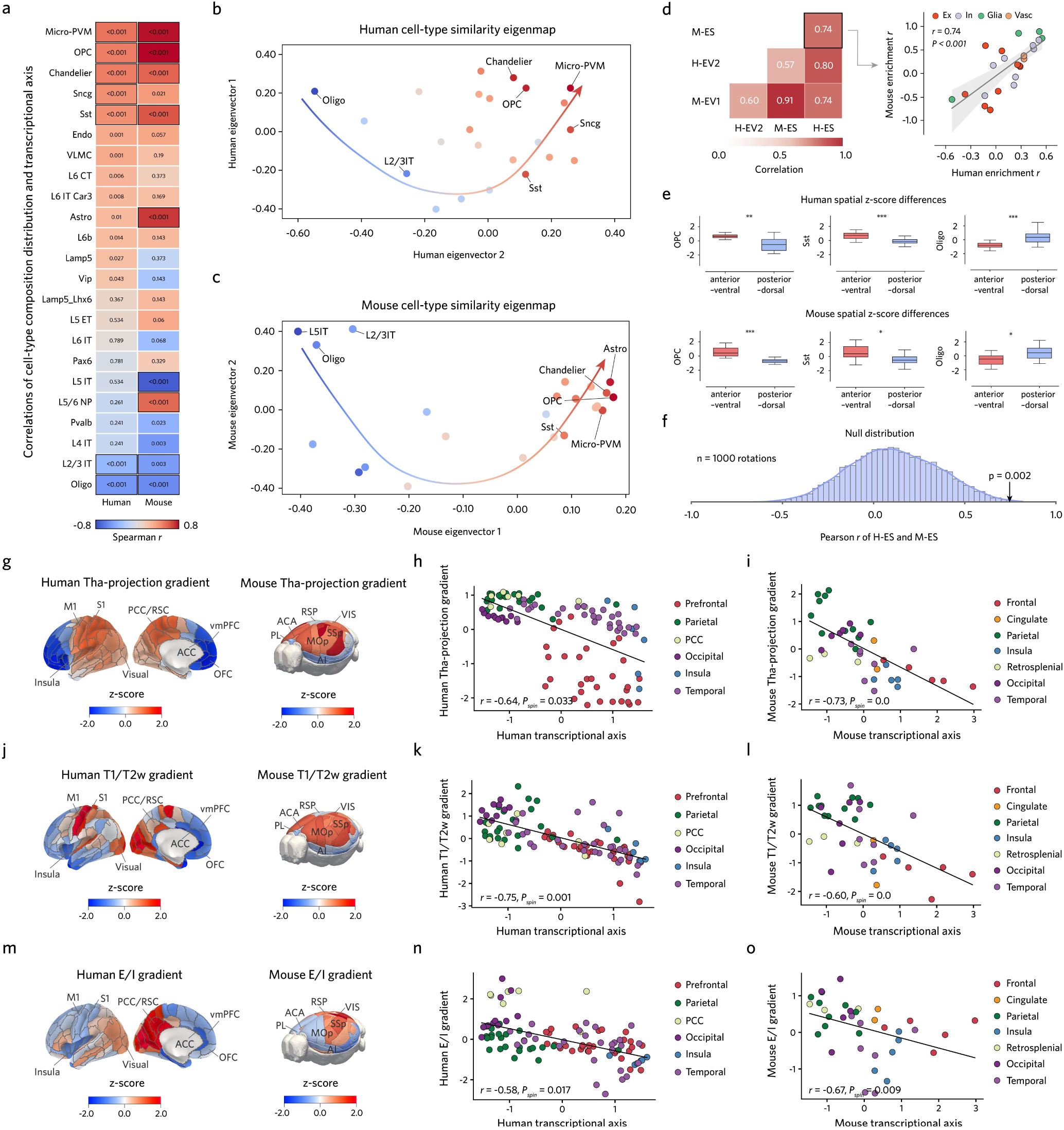
Cortical cell-type composition is organized along the AV–PD axis across species. **(a)** Spearman correlations between the AV–PD transcriptional axis and the cortical distributions of 23 cell types in humans (left) and mice (right). Color indicate Spearman *r* values, and numbers indicate FDR-adjusted *P* values; black borders denote significant spatial correlations (*P_spin_* < 0.05, 1,000 rotations). **(b, c)** Diffusion-map embedding^51^ of cortical cell-type spatial similarity in humans **(b)** and mice **(c)**. Points represent cell types and are colored by their correlations with the AV–PD axis (red, AV-biased; blue, PD-biased). **(d)** Left: pairwise Pearson correlations among cell-type spatial embeddings (H-EV2 and M-EV1) and axis–cell-type association vectors (H-ES and M-ES; *r* = 0.57–0.91, all FDR-adjusted *P* < 0.01, *n* = 23 cell types). Right, scatterplot of the M-ES versus H-ES (Pearson *r* = 0.74, *P* = 5.43×10^−5^); points are colored by cell class and shading denotes the 95% confidence interval of the linear fit. **(e)** AV–PD differences in representative cell-type distributions in humans (OPC, rank-biserial *r* = 0.54, *P* = 1.07 × 10^−^³; Sst, *r* = 0.72, *P* = 2.10 × 10^−^⁵; Oligo, *r* = −0.75, *P* = 1.71 × 10^−^⁵, *n* = 22 AV and 45 PD ROIs) and mice (OPC, *r* = 0.91, *P* = 2.28 × 10^−^⁴; Sst, *r* = 0.59, *P* = 0.02; Oligo, *r* = −0.45, *P* = 0.04, *n* = 14 AV and 17 PD ROIs; two-sided Mann–Whitney *U* test, FDR-corrected). Box: median, 25th–75th percentiles; whiskers, 1.5 × IQR. **(f)** Null distribution of the H-ES–M-ES correspondence (Observed *r* = 0.74, *P_null_* = 0.002, *n* = 1,000 rotations). \**P* < 0.05; \*\**P* < 0.01; \*\*\**P* < 0.001. H-ES and M-ES, human and mouse axis–cell-type association vector; H-EV2 and M-EV1, second human and first mouse eigenvectors of the cell-type spatial embedding, respectively. Ex, excitatory; In, inhibitory; Vasc, vascular.

To determine whether the AV–PD axis represents a principal organizer of cortical cell-type composition, we constructed diffusion-map embeddings^51^ based on spatial similarity among 23 cortical cell types in humans and mice (**Methods**). Cell types were arranged along a continuous cellular axis that closely matched their AV–PD enrichment patterns (**Fig. 3b, c**). The cell-type embedding correlated strongly with the AV–PD gradient association vector in humans (Pearson *r* = 0.80, *P* = 1.38×10^−5^) and mice (*r* = 0.91, *P* = 1.05×10^−8^), with conserved cellular organization across species (*r* = 0.74, *P* = 5.43×10^−5^; **Fig. 3d**). This correspondence remained significant after spatial-spin null testing, cross-dataset replication and cross-species prediction analyses (**Fig. 3e, f**; **Extended Data Fig. 2**; **Supplementary Figs. 3–5; Methods**), demonstrating that the AV–PD axis captures a conserved cellular coordinate rather than a dataset-specific transcriptional pattern.

Beyond cellular composition, the AV–PD coordinate was also reflected in macroscale cortical architecture. The gradient aligned with major cortical phenotypic gradients spanning connectivity, structure and functional organization across species. Along the connectional dimension, the AV–PD axis correlated with thalamocortical projection gradients^34^ (human: Spearman *r* = −0.64, *P_spin_* = 0.033; mouse: *r* = −0.73, *P_spin_* = 0.001; **Fig. 3g–i**), with AV regions characterized by higher-order thalamic input and PD regions by first-order sensory projections. Along the structural dimension, the gradient correlated with cortical myelination, using the T1w/T2w ratio as a proxy^15,35^ (human: Spearman *r* = −0.75, *P_spin_* = 0.001; mouse: *r* = −0.60, *P_spin_* = 0.001; **Fig. 3j–l**), consistent with PD-pole enrichment of myelination-related genes and oligodendrocytes. Along the functional dimension, the AV–PD gradient also correlated with excitation–inhibition organization^36^ (human: Spearman *r* = −0.58, *P_spin_* = 0.017; mouse: *r* = −0.67, *P_spin_* = 0.009; **Fig. 3m–o**), with higher E/I ratios toward the PD pole, paralleling the enrichment of mature neuronal and synaptic signaling programs.

Together, these findings extend the AV–PD axis from a conserved transcriptional gradient to a multi-scale organizational coordinate of the mammalian cortex, linking molecular programs with cell-type composition, thalamocortical connectivity, myelination and excitation–inhibition balance.

### The AV–PD organization is detectable prenatally

Next, we asked whether the adult-defined transcriptional pattern was already spatially organized during development. We held the coordinate fixed and projected adult-derived AV–PD gene weights onto independent developmental datasets^1,25,26,52^ from both species (**Methods**). This approach tested for early spatial expression of the adult-derived gene-weight pattern without re-estimating the axis at each developmental stage.

In humans, adult BrainSpan^1,25,52^ samples reproduced the AV–PD gradient observed in the AHBA^13^ (Spearman *r* = 0.87, *P* = 4.24 × 10^−^⁴). A coarse adult-like pattern was already evident prenatally and was maintained through childhood (prenatal: *r* = 0.62, *P* = 0.04; birth to 13 years: *r* = 0.78, *P* = 4.72 × 10^−3^; **Fig. 4a, b**), consistent with previous observations in the primate brain^19,22^. BrainSpan includes only 11 bulk-dissected neocortical regions, with the AV pole represented by medial and orbital prefrontal cortex alone. These estimates are therefore conservative and do not resolve the gradient continuously across the cortical sheet.

**Figure 4.**
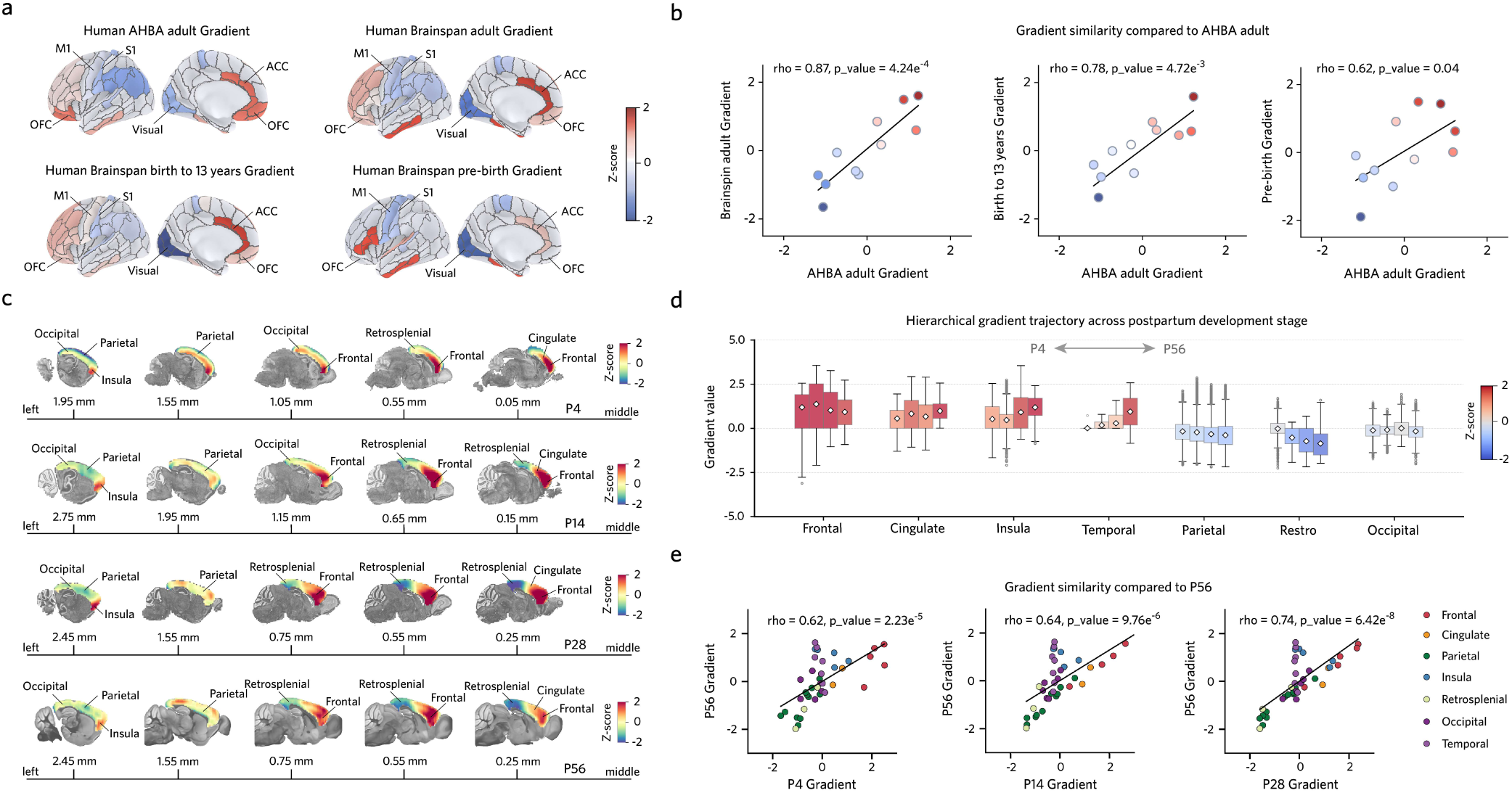
Developmental spatial topology of the AV–PD transcriptional axis. **(a)** Spatial patterns of the AV–PD gradient across human cortical development: adult AHBA^13^ (top-left), BrainSpan^1,52^ adult (top-right), birth to 13 years (bottom-left) and pre-birth (bottom-right). Colors indicate gradient z-scores. (**b**) Correspondence between BrainSpan gradients and the adult AHBA gradient at adult (Spearman *r* = 0.87, *P* = 4.24 × 10^−^⁴), birth to 13 years (*r* = 0.78, *P* = 4.72 × 10^−^³) and pre-birth (*r* = 0.62, *P* = 0.04; *n* =11 ROIs). Points represent cortical regions and are colored by AHBA gradient z-score. (**c**) Sagittal maps of the mouse AV–PD gradient at P4, P14, P28 and P56. Numbers indicate the medial–lateral distance from the midline (mm). Colors indicate gradient z-scores. **(d)** Gradient distributions across seven cortical regions ordered from AV to PD (frontal, cingulate, insula, temporal, parietal, retrosplenial and occipital) at P4, P14, P28 and P56. Diamonds, means. Box: median, 25th–75th percentiles; whiskers, 1.5 × IQR. **(e)** Correspondence between juvenile and adult (P56) mouse gradients (P4, *r*= 0.62, *P* = 2.23 × 10^−^⁵; P14, *r* = 0.64, *P* = 9.76 × 10^−^⁶; P28, *r* = 0.74, *P* = 6.42 × 10^−^⁸; Spearman correlations, *n* = 39 ROIs). Points are colored by lobe. AHBA, Allen Human Brain Atlas; M1, primary motor cortex; S1, primary somatosensory cortex; OFC, orbitofrontal cortex; ACC, anterior cingulate cortex; P, postnatal day.

The mouse data provided complementary spatial and temporal resolution. A whole-cortex spatiotemporal transcriptomic atlas^26^ provided continuous spatial coverage across the cortical sheet and allowed us to trace the adult-derived AV–PD pattern across development. We projected mouse AV–PD gene weights onto this independent atlas, beginning with postnatal stages for which adult cortical areas were resolved in the atlas annotation (P4, P14, P28, and P56; **Methods**). Similar to the human data, the topology of the gradient was consistent across all stages: frontal, cingulate, and insular regions retained AV-pole values, whereas parietal, retrosplenial, and occipital regions retained PD-pole values (**Fig. 4c, d**). The spatial pattern at P4 was already strongly correlated with the adult P56 configuration (Spearman *r* = 0.62, *P* = 2.23 × 10^−^⁵), and the correspondence increased at P14 (*r* = 0.64, *P* = 9.76 × 10^−^⁶) and P28 (*r* = 0.74, *P* = 6.42 × 10^−^⁸; **Fig. 4e**). To extend the analysis to embryonic stages, at which the atlas annotation does not resolve adult cortical areas, we used the sagittal anterior–posterior (A–P) direction as a geometric reference for quantifying the cortical transcriptional gradient (**Methods**). This geometric reference captured the spatial orientation of the postnatal AV–PD axis and provided a common coordinate across prenatal and postnatal stages (E13.5, E15.5, E18.5, P4, P14 and P28; **Fig. 5a**), allowing us to trace the organization back to embryonic development.

**Figure 5.**
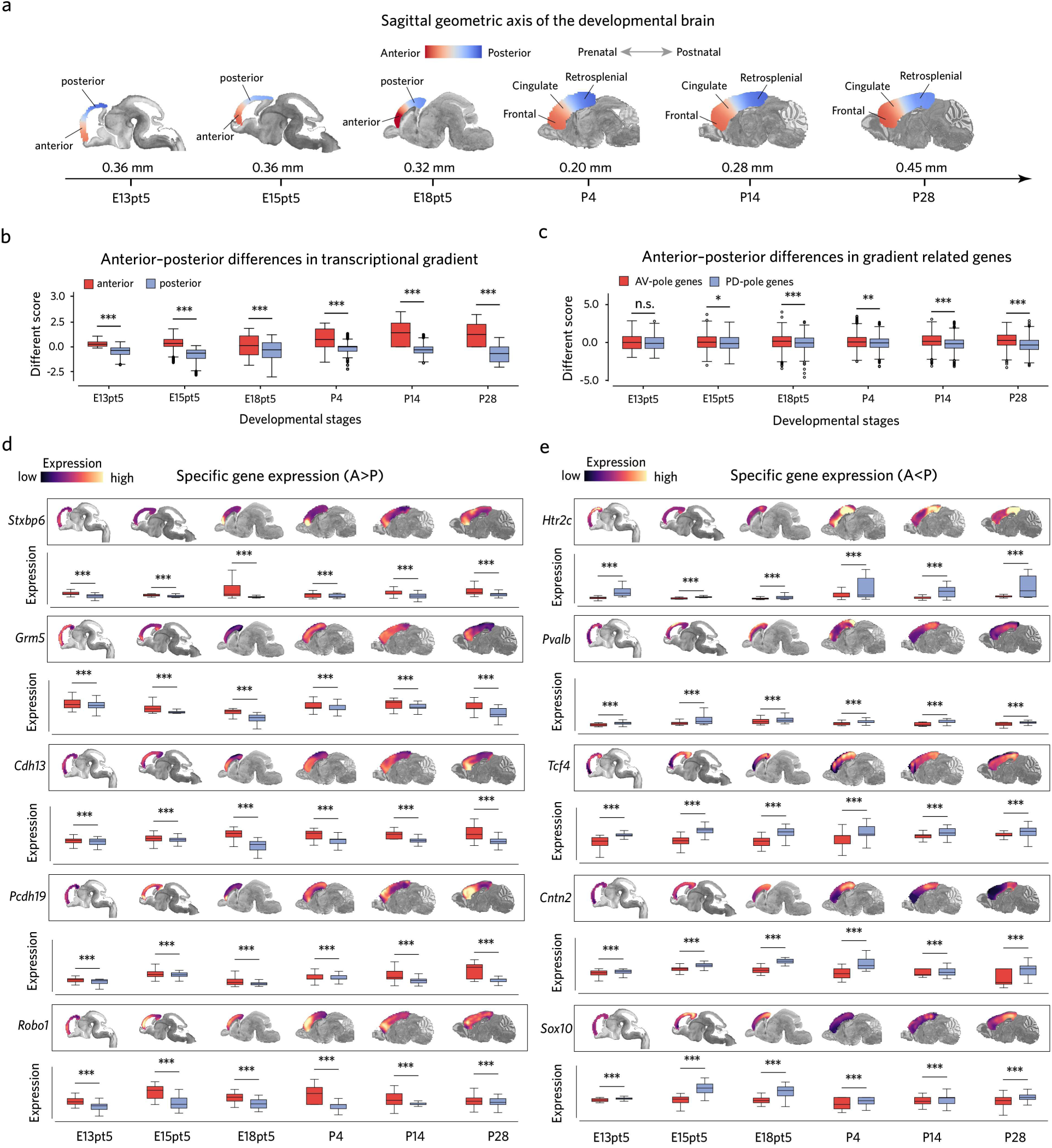
Prenatal spatial organization of the AV–PD transcriptional axis in mice. **(a)** Sagittal slices showing the geometric anterior–posterior reference axis (red, anterior; blue, posterior) used to quantify the AV–PD organization across mouse cortical development. Numbers indicate medial–lateral distance from the midline (mm). **(b)** Anterior–posterior differences in transcriptional gradient z-scores across developmental stages. (rank-biserial *r* from E13.5 to P28: 0.91, 0.82, 0.37, 0.60, 0.90 and 0.92; all FDR-adjusted *P* < 0.001; two-sided Mann–Whitney *U* test). Box: median, 25th–75th percentiles; whiskers, 1.5 × IQR. **(c)** Anterior–posterior expression differentials for AV-biased (*n* = 566) and PD-biased (*n* = 591) genes (rank-biserial *r* from E13.5 to P28: 0.04, 0.18, 0.12, 0.19, 0.24 and 0.35; *P* = 0.22, 0.023, 8.25 × 10^−^⁴, 6.0 × 10^−^⁴, 7.08 × 10^−^¹² and 1.89 × 10^−^²², respectively; two-sided Mann–Whitney *U* test; FDR corrected). Box: median, 25th–75th percentiles; whiskers, 1.5 × IQR. **(d, e)** Developmental spatial expression of representative AV-pole genes (**d**; *Stxbp6, Grm5, Cdh13, Pcdh19* and *Robo1*) and PD-pole genes (**e**; *Htr2c, Pvalb, Tcf4, Cntn2* and *Hivep2*). Sagittal maps show expression levels (z-score) and box plots compare anterior and posterior regions across six stages (all *P* < 0.001, FDR-corrected; two-sided Mann–Whitney *U* test). Box: median, 25th–75th percentiles; whiskers, 1.5 × IQR. n.s., not significant; \**P* < 0.05; \*\**P* < 0.01; \*\*\**P* < 0.001. E, embryonic day; P, postnatal day.

We first tested whether the projected gradient was spatially organized along this axis at each stage. A significant difference between the AV and PD poles was already detectable at E13.5 and persisted across all subsequent stages (E13.5: rank-biserial *r* = 0.91; E15.5: *r* = 0.82; E18.5: *r* = 0.37; P4: *r* = 0.60; P14: *r* = 0.9; P28: *r* = 0.92; all *P* < 0.001; two-sided Mann–Whitney U tests; **Fig. 5b**). However, a detectable aggregate gradient does not require broad polarization of the adult-defined AV- and PD-pole gene sets. We therefore next asked when the two gene sets acquired their mature spatial polarity. For each gene, we calculated the expression differential along the geometric axis as anterior minus posterior expression and compared the distributions of the adult-defined AV- and PD-pole gene sets at each stage (**Methods**). The two distributions were not significantly separated at E13.5 (rank-biserial *r* = 0.04, *P* = 0.22). From E15.5 onward, however, AV-pole genes showed progressively higher differential scores than PD-pole genes (E15.5: *r* = 0.18, *P* = 0.023; E18.5: *r* = 0.12, *P* = 8.25 × 10^−^⁴; P4: *r* = 0.19, *P* = 6.0 × 10^−^⁴; P14: *r* = 0.24, *P* = 7.08 × 10^−^¹²; P28: *r* = 0.35, *P* = 1.89 × 10^−^²²; two-sided Mann–Whitney U tests; **Fig. 5c**). Thus, a weighted transcriptional difference between the future poles was already detectable at E13.5, whereas broader polarization of the adult gene sets emerged from E15.5 onward.

We also identified genes that maintained stable AV- or PD-pole spatial polarity across the developmental stages examined (**Supplementary Table 1**). Stable AV-pole genes were associated with cortical regionalization, cell adhesion and axon guidance, such as *Pcdh19*^53^, *Cdh13*^54^ and *Robo1*^55^ (**Fig. 5d**). Stable PD-pole genes reflected mature signaling, myelination and circuit maintenance, such as *Pvalb*^56^, *Cntn2*^57^ and *Sox10*^58^ (**Fig. 5e**). These representative genes recapitulated the broader functional distinction between cortical construction at the AV pole and mature signaling and myelination at the PD pole.

These findings support a prenatal origin of the AV–PD coordinate, which was progressively refined and maintained into adulthood.

### Cortical developmental programs unfold along the AV–PD axis

The progressive spatial maturation of AV–PD organization prompted us to ask whether this stable spatial coordinate also structured the temporal unfolding of cortical gene-expression programs. For each cortical region, we calculated the developmental rate of expression change for each gene and decomposed the resulting region-by-gene slope matrix (**Fig. 6a**; **Methods**). The principal components of developmental change were strongly correlated with the AV–PD axis in both species (human: Spearman *r* = 0.84, *P_spin_* = 0.0013; mouse: *r* = 0.83, *P_spin_* = 0.04; **Fig. 6a**). Conversely, weighting adult cortical expression by the developmental-slope loadings reproduced the AV–PD spatial gradient (**Fig. 6b**). These analyses linked the adult spatial axis to the dominant regional pattern of developmental expression change. We next resolved this relationship along its gene-program and regional dimensions.

**Figure 6.**
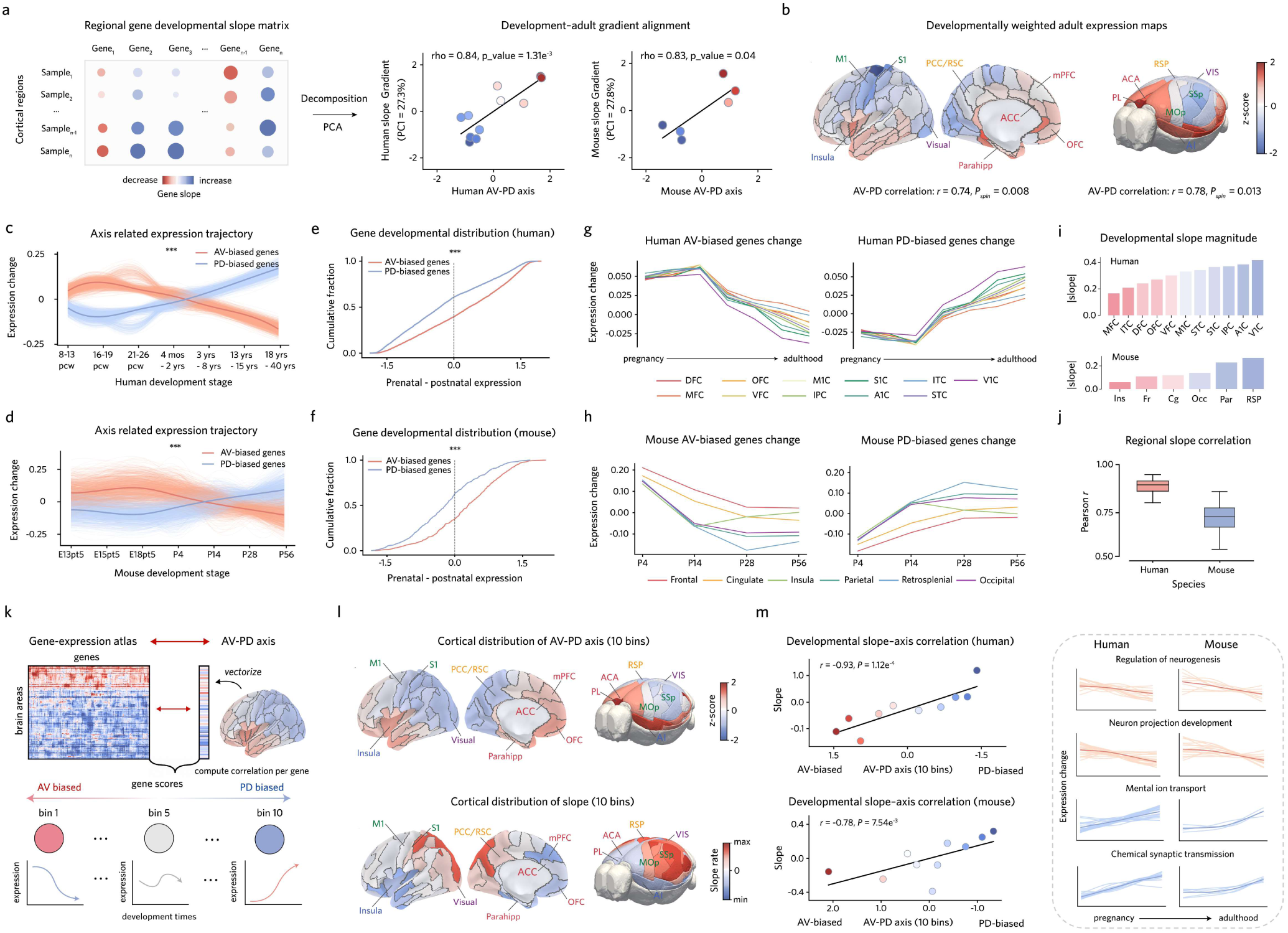
The AV–PD transcriptional axis links to conserved developmental timing across species. **(a**) PCA of the developmental slope matrix. PC1 correlated with the adult AV–PD axis in humans (Spearman *r* = 0.84, *P* = 0.0013, *n* = 11 ROIs) and mice (*r* = 0.83, *P* = 0.04, *n* = 6 ROIs). **(b)** Adult-expression-weighted loadings recapitulated the AV–PD organization in humans (*r* = 0.74, *P_spin_* = 0.008) and mice (*r* = 0.78, *P_spin_* = 0.013). **(c, d)** Developmental trajectories of AV- and PD-biased genes in humans (**c**; *n* = 5,913 and 5,709) and mice (**d**; *n* = 566 and 591). Light lines, 1,000 bootstrap trajectories; bold lines, means. Trajectories differed in both species (human, *D_traj_* = 5.20; mouse, *D_traj_* = 3.67; both *P* = 0.001, 1,000 permutations; *D_traj_*: the summed squared distance between the mean trajectories). **(e, f)** Cumulative distributions of prenatal-to-postnatal expression bias in humans (**e**) and mice (**f**), showing greater prenatal expression of AV-biased genes (rank-biserial *r* = 0.25 and 0.34, respectively; both *P* < 0.001, two-sided Mann–Whitney *U* tests). Dashed lines indicate zero bias. **(g, h)** Regional developmental changes of AV- and PD-biased genes in humans (**g**) and mice (**h**). **(i)** Regional-slope magnitudes across cortex along the AV–PD axis. **(j)** Pearson correlations of regional developmental slope vectors, indicating concordant developmental directions of genes across regions. Box: median, 25th–75th percentiles; whiskers, 1.5 × IQR. (**k)** Genes ranked into ten bins from AV- to PD-biased expression. (**l)** Adult expression distributions of bins (top) and their developmental slopes (bottom). (**m)** Developmental slope versus adult expression across bins in humans (Spearman *r* = −0.93; *P* = 1.12×10^−4^) and mice (Spearman *r* = −0.78; *P* = 7.54×10^−3^). Right, trajectories of representative AV-pole (orange) and PD-pole (blue) enriched modules. Light lines, genes; dark lines, module means. Brain region abbreviations in **Supplementary Table 3.** \*\*\**P* < 0.001. pcw, post-conception week; mos, months; yrs, years; E, embryonic day; P, postnatal day.

At the gene-program level, AV- and PD-biased genes followed opposing developmental trajectories relative to the genome-wide mean (**Methods**). AV-biased genes peaked during the prenatal neurodevelopmental window (human: 16–26 post-conceptional weeks; mouse: E15.5–E18.5), whereas PD-biased genes remained relatively low. Their trajectories subsequently crossed and reversed polarity, and this configuration was maintained into adulthood (human: 18–40 years; mouse: P56). The developmental courses of the two gene sets differed significantly in both species (human: *D_traj_* = 5.20, *P_permutation_* = 0.001; mouse: *D_traj_* = 3.67, *P_permutation_* = 0.001; 1,000 label permutations; **Fig. 6c, d**; **Methods**). Consistent with the opposing temporal profiles of the two gene sets, a group-level distributional analysis showed that AV-biased genes shifted toward prenatal expression and PD-biased genes toward postnatal expression in both species (**Fig. 6e, f**). The same temporal polarity was also evident at the single-gene level. Prenatal-to-postnatal differential expression correlated positively with AV–PD gene weights in both humans (Pearson *r* = 0.17, *P* = 1.69 × 10^−^⁶⁵) and mice (*r* = 0.22, *P* = 9.85 × 10^−^¹⁵), with AV-biased genes showing higher prenatal expression and PD-biased genes showing higher postnatal expression (**Extended Data Fig. 4a, b**).

At the regional level, these opposing expression changes occurred throughout the cortex but differed systematically in magnitude along the AV–PD axis. In humans, anterior and prefrontal regions (DFC, MFC, and OFC) showed the shallowest decline in AV-biased genes and retained the highest adult expression, whereas posterior sensory regions (V1C and S1C) showed the steepest decline and lowest adult expression; PD-biased genes showed the opposite pattern (**Fig. 6g, i**; top). In mice, insular, frontal, and cingulate regions changed least in both directions, whereas parietal and occipital regions changed most (**Fig. 6h, i**; bottom). Developmental slope vectors nevertheless remained highly correlated across regions in both species (**Fig. 6j**), indicating that cortical regions shared the direction of developmental change but differed in its magnitude.

This relationship further extended across the full gene-expression continuum. We ranked genes by their adult spatial correlation with the AV–PD transcriptional axis, divided them into 10 bins from AV-biased to PD-biased expression, and calculated the mean developmental slope of each bin (**Fig. 6k**). The spatial-bias bins paralleled the regional organization of developmental slopes in both species (**Fig. 6l**), and adult spatial bias was negatively associated with developmental slope across bins (human: Spearman *r* = −0.93, *P* = 1.12 × 10^−^⁴; mouse: *r* = −0.78, *P* = 7.54 × 10^−^³; **Fig. 6m**, left).

Additional evidence also supports this temporal unfolding pattern. From GO enrichment (**Fig. 2f**), AV-pole genes in neurogenesis and neuronal projection modules were highly expressed prenatally and downregulated after birth, whereas PD-pole genes involved in metal-ion transport and chemical synaptic transmission were progressively upregulated into adulthood (**Fig. 6m**, right). Canonical developmental transitions showed the same polarity. Based on established cell-type markers and developmental transcriptomic atlases^1,59^, adult AV-pole expression encompassed programs associated with neural and glial progenitors (*HES5, SOX2*, *ID4*, and *PDGFRA*), immature neurons (*TUBB2A* and *TUBB3*) and early synaptic function (*GRIN3A*, *GABRA5*, *GABRB3*, *GLRA2*). By contrast, the PD pole encompassed neuronal identity and maturation (*RORB*, *EMX1*, *POU3F2*, and *INA*), myelination (*MBP*) and mature synaptic machinery (*GRIN2A*, *GABRA1*, *GLRB*, *PVALB* and *KCNC1*) (**Extended Data Fig. 3**). The *GRIN3A*-to-*GRIN2A* NMDA receptor switch^60^ and the *GABRA5*/*GABRB3*-to-*GABRA1* receptor switch^61^ were also resolved across opposite poles, with the immature components at the AV pole and the mature components at the PD pole. The temporal polarity also extended to multiple regulatory layers. AV-pole genes showed a greater prenatal shift in cis-eQTL effect size, whereas PD-pole genes showed larger developmental gains in active enhancer and transcription-start-site states and greater losses of Polycomb-repressed states (**Extended Data Fig. 4c, d**). The two poles were also associated with distinct transcriptional regulators: AV-pole genes were enriched for neurogenic Wnt/bHLH, SOX9 and REST/NRSF modules^62–64^, whereas PD-pole genes were enriched for activity-dependent and mature-neuronal CREB, MEF2 and ERRα modules^65–67^ (**Extended Data Fig. 4e**).

Together, these analyses show that cortical regions follow a broadly shared direction of developmental transcriptional change but differ systematically in its temporal unfolding along the AV–PD axis. The mature spatial organization of the cortex therefore retains a graded spatial imprint of developmental timing: the AV pole preserves more developmental features related to cortical construction, whereas features associated with mature neuronal function progressively dominate toward the PD pole.

### Autism-associated perturbations converge along the conserved AV–PD axis

We next asked whether the AV–PD axis is associated with neurodevelopmental susceptibility. Several genes polarized along the axis are established risk genes for autism spectrum disorder (ASD), intellectual disability (ID), or schizophrenia (SCZ), including *PCDH19*, *GABRB3*, and *GABRA5* at the AV pole and *GRIN2A*, *GABRA1*, *TCF4*, *POU3F2*, and *RORB* at the PD pole^47,68–70^. Disease-associated cell types are similarly distributed along the axis: Sst and chandelier interneurons at the AV pole and supragranular L2/3 IT projection neurons at the PD pole show prominent transcriptomic, synaptic, and connectivity alterations in ASD and SCZ^71–73^. Human studies have associated this axis with disease-risk gene expression^22^,cortical volume alterations^22^, and ASD-related shifts in transcriptional identity^46^. However, these population-level associations ignore heterogeneous genetic background. We therefore tested whether genetically diverse ASD mutations converge on the AV–PD axis in developmental transcription and cortical anatomy.

We first analyzed whole-cortex single-nucleus transcriptomic data^37^ from 11 ASD mouse models from E14.5 to P14. For each gene, we calculated its developmental expression slope separately in WT and ASD mice. Because these data lacked spatial coordinates, we used each gene’s adult AV–PD expression bias as a reference and divided genes into 10 bins ranging from AV-biased to PD-biased expression (**Methods**). This strategy linked developmental dysregulation to the adult AV–PD cortical coordinate in the absence of direct measurements of regional gene expression. ASD-associated dysregulation was quantified by comparing developmental slopes in bins between ASD and WT mice (**Fig. 7a**; **Methods**). In WT mice, developmental slopes varied continuously across the bins: AV-biased genes generally showed negative slopes, whereas slopes became progressively more positive toward PD-biased genes, recapitulating the developmental gradient described above. ASD mice retained this overall ordering but showed a systematic shift toward more negative slopes (**Fig. 7b**). The ASD–WT difference became increasingly stronger toward the PD-biased end and was strongly correlated with adult AV–PD position (Spearman *r* = −0.85, *P* = 1.78 × 10^−^³; **Fig. 7c**). This shift was broadly reproduced across genetic models. Ten of the 11 ASD genotypes showed a negative mean slope shift relative to WT, and the bin-wise pattern was significantly correlated with the adult AV–PD gradient in eight genotypes (**Fig. 7d**). Genetically heterogeneous ASD models therefore shared an axis-organized developmental displacement, although its magnitude varied across genotypes.

**Figure 7.**
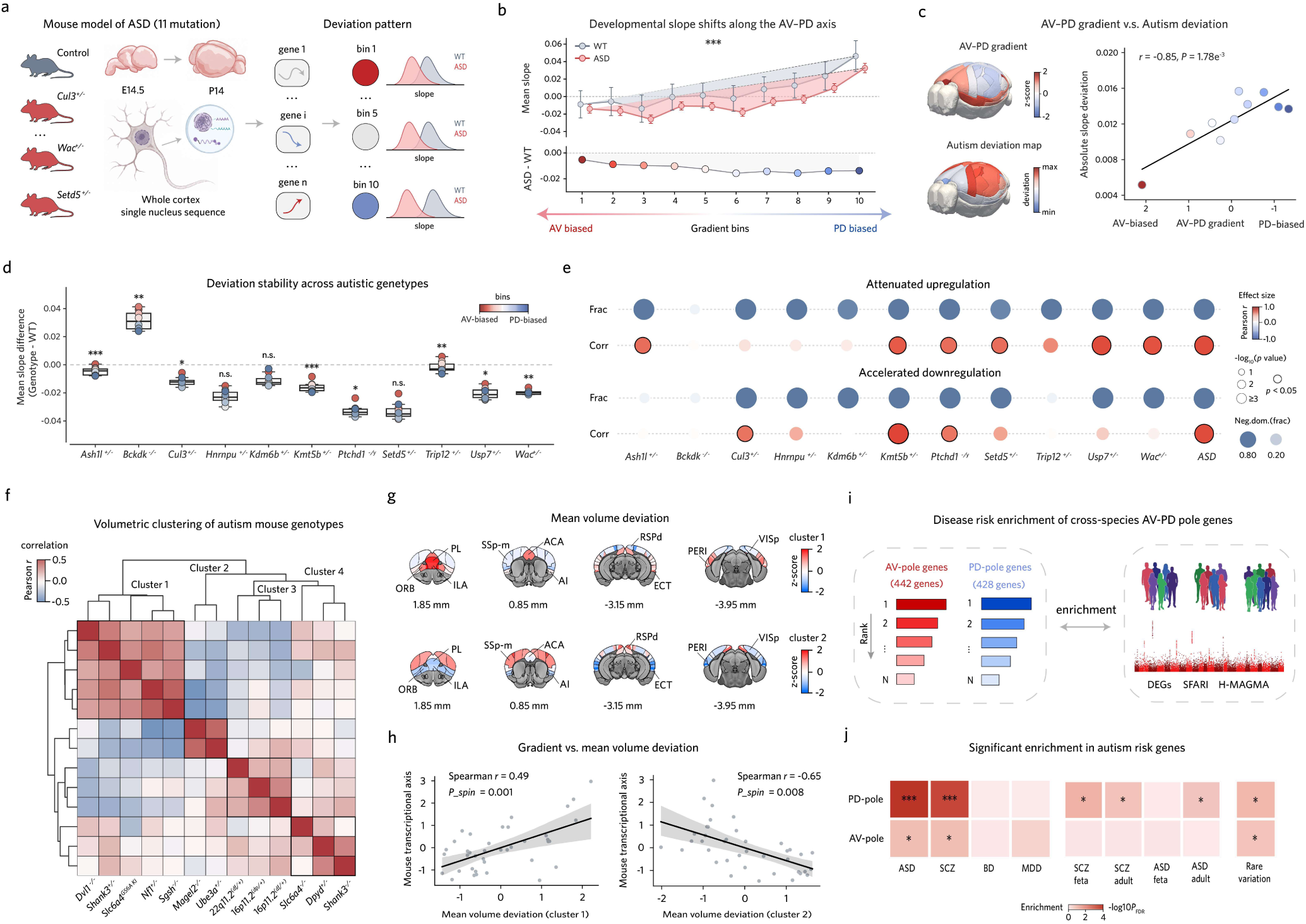
Autism-related molecular and anatomical alterations converge along the AV–PD axis. **(a)** Schematic of developmental transcriptomic analysis across 11 autism mouse genotypes. Gene-wise developmental slopes were summarized across ten AV–PD bins and compared with wild type (WT). **(b)** Developmental slopes across bins in WT and autism models (top) and ASD– WT differences (bottom). **(c)** Correspondence between the AV–PD gradient and autism-related developmental deviation. Absolute slope deviation increased toward the PD-biased end (Spearman *r* = −0.85, *P* = 1.78 × 10^−^³). **(d)** Developmental slope deviations across autism genotypes. Points are colored by AV–PD bin. Box: median, 25th–75th percentiles; whiskers, 1.5 × IQR. **(e)** Autism-related negative slope shifts were decomposed into attenuated upregulation and accelerated downregulation effects across AV–PD bins. For Corr, bubble color denotes similarity between affected-bin patterns and adult AV–PD bin expression, bubble size indicates significance, and black borders mark significant correlations; for Frac, bubble size represents the fraction of affected bins. **(f)** Hierarchical clustering of volumetric deviations across autism genotypes, identifying four clusters. **(g)** Mean regional volume-deviation maps for clusters 1 and 2. Colors indicate z-scores. **(h)** Cluster-level volume deviations versus the AV–PD axis (cluster 1: Spearman *r* = 0.49, *P_spin_* = 0.001; cluster 2: *r* = −0.65, *P_spin_* = 0.008, *n* = 39 ROIs). Shading denotes 95% confidence intervals. **(i, j)** Disease-risk enrichment analysis of cross-species AV- and PD-pole genes (**i**) and enrichment across neuropsychiatric risk categories (**j**). Colors indicate -log10(*P_FDR_)*. *\*P <* 0.05; for enrichment analyses, *\*P <* 0.1; \*\**P* < 0.01; \*\*\**P* < 0.001. DEG, differentially expressed genes; ASD, autism spectrum disorder; SCZ, schizophrenia; BD, bipolar disorder; MDD, major depressive disorder; H-MAGMA, Hi-C–coupled MAGMA; SFARI, Simons Foundation Autism Research Initiative.

We next separated genes according to the direction of their developmental slopes in WT mice (downregulated or upregulated) and, within each AV–PD bin, examined how the two gene sets were altered in ASD mice. The negative shift in ASD arose through two changes: genes normally downregulated during development declined more rapidly, whereas genes normally upregulated showed an attenuated increase. Most genotypes exhibited at least one alteration (**Fig. 7e**; **Method**), indicating that distinct mutations can produce a similarly organized displacement through different pathological mechanisms during development.

We next asked whether directly measured structural alterations in ASD mouse models were spatially organized along the same coordinate. Hierarchical clustering of whole-cortex volumetric deviation profiles across 13 ASD-related genotypes^38^ identified four clusters (**Methods**), two of which showed opposing AV– PD patterns (**Fig. 7f**). Cluster 1 (*Dvl1^⁻/⁻^, Shank3^⁺/⁻^*, *Slc6a4^G56A KI^*, *Nf1^⁺/⁻^*, and *Sgsh^⁻/⁻^*^)^ showed relative volume expansion in AV-pole regions (ORB, PL, ILA, AI, and ECT) and contraction in PD-pole regions (VIS, SMA, and RSP). Cluster 2 (*Ube3a^⁺/⁻^* and Magel2^⁻/⁻^) showed the inverse pattern despite an overall reduction in cortical volume (**Supplementary Table 2**). The two profiles were aligned with the AV–PD axis in opposite directions (Cluster 1: Spearman *r* = 0.49, *P_spin_* = 0.001; Cluster 2: *r* = −0.65, *P_spin_* = 0.008; **Fig. 7g, h**). Thus, ASD-related anatomical alterations exist a form of AV-PD axis constrained heterogeneity.

As complementary molecular evidence, we tested conserved AV–PD genes for enrichment across H-MAGMA risk genes^39,40^, disease-associated differentially expressed genes^41–46^ (DEGs), and SFARI rare-variant ASD genes^47^ (**Fig. 7i**; **Methods**). Enrichment for ASD- and SCZ-associated genes was detected at both poles but was stronger at the PD pole (**Fig. 7j**). PD-pole genes were enriched across disease-associated DEGs, fetal and adult SCZ H-MAGMA genes, and SFARI genes, whereas AV-pole genes showed more modest enrichment among disease-associated DEGs and SFARI genes.

Together, these findings identify the conserved AV–PD axis as a shared coordinate of autism-associated cortical alterations. Genetically diverse mutations converged on this coordinate in developmental transcription and cortical anatomy, although individual genotypes differed in the magnitude, direction and developmental mechanism of their alterations.

## Discussion

Despite extensive molecular and structural divergence between the human and mouse cortex, it remains unclear whether their molecular and cellular features are assembled into a conserved, cortex-wide program of organization and development. Here, we identified an AV–PD transcriptional axis that was independently recovered in human and mouse cortical transcriptomes. This axis provides a common spatial coordinate for homologous gene expression, cortical cell-type proportions and macroscale phenotypes across species. Its adult-like configuration was already detectable prenatally and was progressively refined without reorientation. Along this relatively stable spatial coordinate, AV-biased genes retained early developmental signatures and declined across development, whereas PD-biased genes progressively engaged mature neuronal and synaptic programs. Genetically diverse autism mouse models preserved this overall developmental ordering but showed a shared negative displacement that increased toward the PD pole. Together, these findings distinguish the AV–PD axis as the cortical readout of a conserved, gene-defined developmental program linking spatial organization, developmental timing and autism-associated vulnerability.

Previous studies have frequently interpreted the principal transcriptional gradient of the human and macaque cortex in relation to the sensorimotor–association (S–A) hierarchy, which extends from primary sensorimotor regions to transmodal association cortex^13,15,16^. Our results indicate that the canonical S–A hierarchy and the dominant cortical transcriptional gradient should not necessarily be treated as equivalent. The AV–PD axis extends primarily from limbic and paralimbic territories—including cingulate, insular, orbitofrontal and parahippocampal cortices—to sensorimotor and visual regions. Thus, although the AV– PD and S–A axes share a primary sensory pole, they diverge at their opposing poles. This distinction is consistent with recent transcriptomic decomposition studies identifying multiple partly independent components in the human cortex^22–24^. The principal component C1 exhibits a limbic-to-sensory-visual organization that corresponds closely to the AV–PD axis, whereas C3 more closely resembles the canonical S–A hierarchy and includes the lateral prefrontal cortex. Recent work has also described a related primary sensory–allocortical (Pr–Al) molecular organization in marmoset cortex^19^, further suggesting that transcriptional differentiation between limbic or allocortical and primary sensory territories may represent a broader feature of cortical organization. The distinction between these axes becomes particularly important in mammalian comparisons. Some of the transmodal association regions anchoring the primate S–A hierarchy, particularly granular lateral prefrontal and inferior parietal cortex, lack clear homologues in mouse. By contrast, the limbic, sensorimotor and visual territories defining the AV–PD axis have more readily identifiable counterparts.

Whether a conserved cortical axis spanning transcriptomic, cellular, and mesoscale phenotypes and linked to developmental dynamics exists in both humans and mice has remained poorly understood. Across four independent whole-cortex mouse transcriptomic datasets^26,28–30^, the AV–PD gradient was consistently recovered through unsupervised decomposition and corresponded significantly to the independently derived human axis. Homologous genes also made corresponding, although quantitatively modest, contributions to the human and mouse gradients, suggesting that a common spatial organization can be maintained despite evolutionary reconfiguration of its individual molecular components. Across species, AV regions were enriched for cellular signatures related to OPCs, microglia/perivascular macrophages, and specific inhibitory interneuron classes including Sst, Sncg, and chandelier cells, and were associated with higher-order thalamic input and lower T1w/T2w values. In contrast, PD regions were enriched for mature oligodendrocytes and intratelencephalic excitatory projection neurons, and were associated with first-order sensory input, higher T1w/T2w values, and higher E/I ratios. This shared polarity supports conservation of a multiscale organizational coordinate rather than resemblance between transcriptional maps alone.

Developmentally, the adult-derived AV–PD orientation was detectable during embryonic development, whereas broader polarization of its constituent gene programs emerged progressively thereafter. The axis is neither generated de novo in adulthood nor developmentally static: its broad spatial orientation emerges early, while the molecular programs defining its mature polarity continue to unfold across development. AV-pole genes were preferentially expressed prenatally and subsequently declined, with enrichment for neurogenesis, axon development, neuronal projection and synaptic organization. PD-pole genes instead increased toward adulthood and were enriched for ion transport, synaptic transmission, myelination and mature neuronal function. Cortical regions shared these overall directions of change but differed continuously in their magnitude, such that the adult spatial position of a gene or region predicted its developmental trajectory. The mature AV–PD gradient therefore retains a graded spatial imprint of developmental timing rather than representing a simple division between immature and mature cortical regions. Developmental shifts in receptor-subunit expression—including GRIN3A-to-GRIN2A among NMDA receptor genes and GABRA5/GABRB3-to-GABRA1 among GABA receptor genes—were resolved across opposite poles of the axis, and the same temporal polarity extended to cis-regulatory effects, chromatin remodeling and transcriptional-regulator associations. These observations suggest that AV–PD organization is embedded in coordinated developmental regulation rather than arising solely from regional variation in adult gene expression.

The AV–PD coordinate also provides a framework for interpreting heterogeneous autism-associated effects. Across genetically diverse autism mouse models, the normal transition from negative developmental slopes among AV-biased genes to increasingly positive slopes among PD-biased genes were retained but displaced in a negative direction. This displacement arose through accelerated downregulation of normally decreasing programs, attenuated upregulation of normally increasing programs, or both, and became more pronounced toward the PD pole. The greater PD-pole displacement is consistent with impaired engagement of later neuronal, synaptic and ion-channel programs^37^, whereas the simultaneous acceleration of developmental downregulation indicates that these abnormalities cannot be reduced to a general developmental delay. Instead, distinct mutations appear to disrupt different components of the same spatially organized developmental program. Structural MRI phenotypes were also organized along the AV– PD coordinate, although different genotype clusters exhibited opposing anatomical patterns. Together with the enrichment of ASD- and schizophrenia-associated genes at both poles, particularly the PD pole, these results identify a form of axis-constrained heterogeneity: diverse genetic perturbations converge on a shared cortical coordinate while differing in the direction, magnitude and developmental mechanism of their effects. The AV–PD axis may define a coordinate of cortical vulnerability rather than a single pathological pathway or a uniform anatomical signature of ASD.

Several limitations warrant further consideration. Human cell-type maps partly relied on cross-species inference from macaque data^31,50^, and projection of adult-derived gene weights demonstrates early developmental alignment but not reveal when or how the axis first originates. Moreover, the observed associations do not establish causality, and conservation of AV–PD organization also does not imply identical molecular implementation across species. The recently proposed MIND (Multinodal Induction– Exclusion in Network Development) model provides one potential mechanistic framework in which competing transcriptional programs emerging from frontotemporal and primary sensorimotor territories contribute to cortical axis formation^74^. Our findings are also consistent with a broader cortical developmental model in which intrinsic molecular patterning establishes early differentiation between limbic-related and sensory cortical territories, while thalamocortical input and activity-dependent processes progressively refine their molecular and cellular organization^75–77^. Under this model, the adult AV–PD axis preserves the spatial consequences of developmental programs that unfold at different rates across the cortex. However, the present analyses establish spatial and temporal correspondence rather than the causal interactions that generate the axis. In the future, spatially resolved developmental data and experimental perturbations of candidate regulatory programs will be needed to address these questions. Nevertheless, by independently recovering the AV–PD axis in humans and mice and linking it to homologous genes, cell types, macroscale phenotypes and developmental timing, our study provides evidence that a core dimension of mammalian cortical organization is built upon an evolutionarily conserved developmental program and constitutes a shared coordinate of vulnerability to neurodevelopmental disruption.

## Extended data figure titles and legends

**Extended Data Figure 1.**
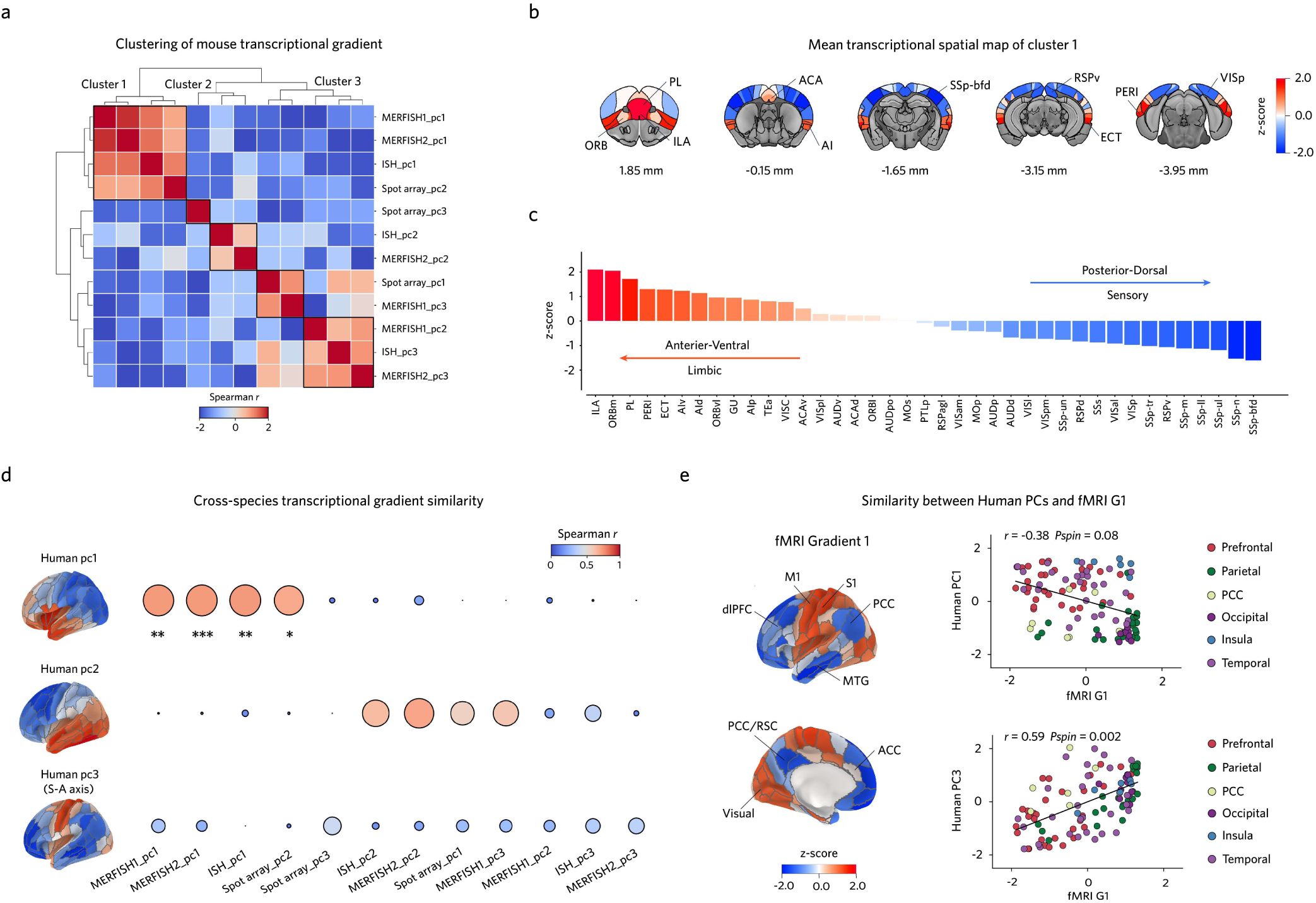
A conserved cross-species cortical transcriptional gradient specific to the AV–PD axis. **(a)** Hierarchical clustering of pairwise Spearman correlations among PC1–PC3 from four independent mouse cortical transcriptomic datasets (ISH^27^, Spot array^28^, MERFISH-1^29^, MERFISH-2^30^). Cluster 1 contains components from all four datasets. **(b)** Mean spatial map of Cluster 1. Numbers indicate anterior–posterior positions relative to bregma (mm); colors indicate z-scores. **(c)** Mean Cluster 1 z-scores across cortical regions ordered from anterior–ventral/limbic to posterior–dorsal/sensory cortex. **(d)** Cross-species similarity between the first three human transcriptional PCs and the 12 mouse components ordered as in **a**. Bubble size and color indicate Spearman correlation strength. Human PC1 selectively correlated with the four Cluster 1 components (all *P_spin_* < 0.05). **(e)** Relationship of human transcriptional PCs with the cortical fMRI gradient 1. Human PC1 showed no significant correspondence with fMRI G1^17^ (Spearman *r* = −0.38, *P_spin_* = 0.08), whereas human PC3, corresponding to the sensorimotor–association axis, was significantly correlated with fMRI G1 (*r* = 0.59, *P_spin_* = 0.002). Points are colored by cortical lobe. \**P* < 0.05; \*\**P* < 0.01; \*\*\**P* < 0.001. PC, principal component; ISH, in situ hybridization; MERFISH, multiplexed error-robust fluorescence in situ hybridization; S–A, sensorimotor–association. Brain region abbreviations in **Supplementary Table 3.**

**Extended Data Figure 2.**
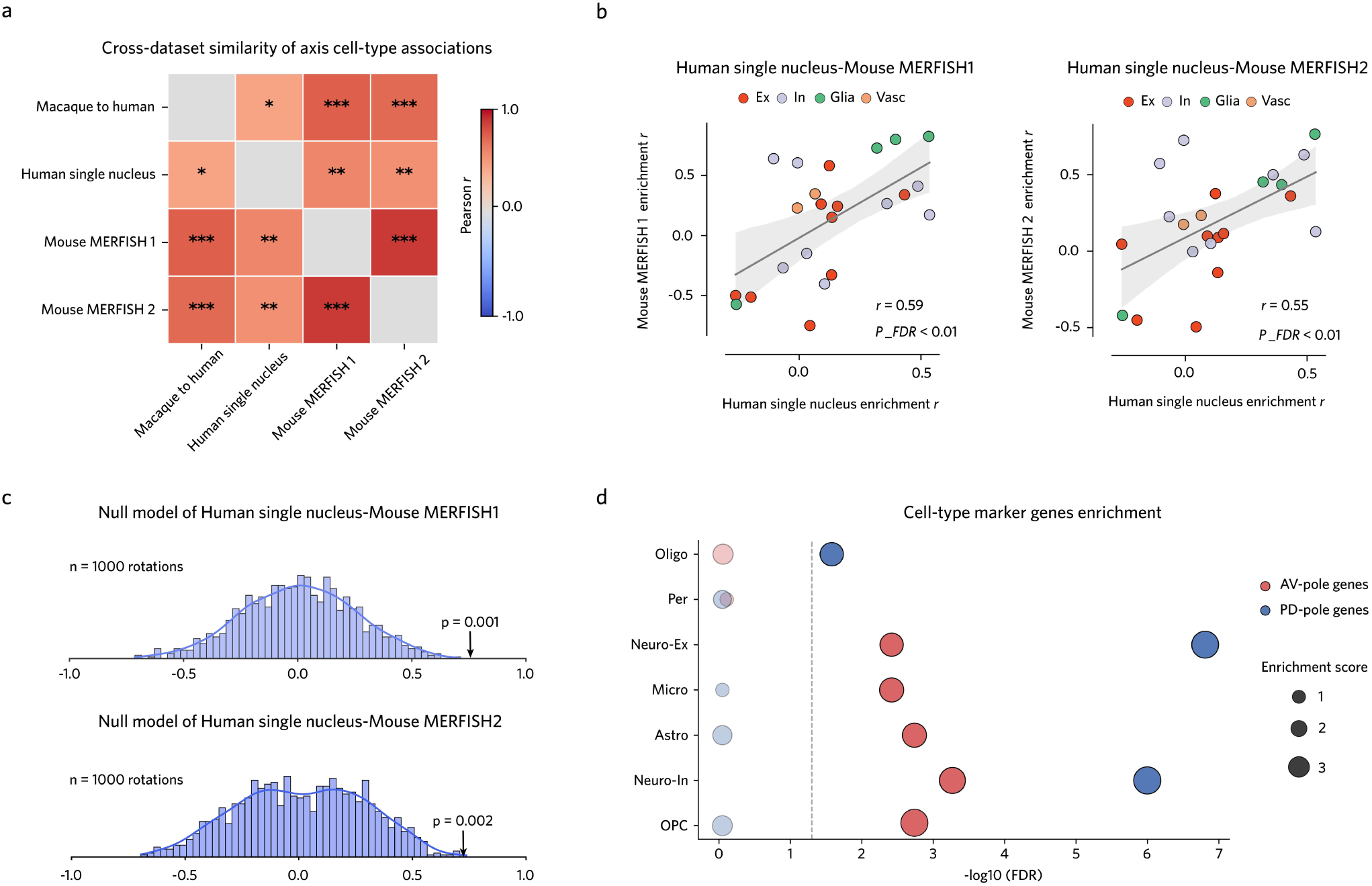
Cross-dataset validation of AV–PD axis–cell-type associations. **(a)** Cross-dataset similarity of AV–PD axis–cell-type association vectors derived from four independent datasets: macaque-to-human spatial inference^31,50^, human single-nucleus RNA-seq^32^, and two mouse MERFISH datasets (MERFISH 1^29^, MERFISH 2^30^). All pairwise correlations were significant after FDR correction (mean *r* = 0.65; *n* = 23 cell types). Colors indicate Pearson *r*. **(b)** Cross-species correspondence of axis– cell-type associations between human single-nucleus data and mouse MERFISH-1 (left, Pearson *r* = 0.59, *P_FDR_* = 4.62 × 10^−^³) or MERFISH 2 (right, Pearson *r* = 0.55, *P_FDR_* = 8.52 × 10^−^³). Points represent cell types and are colored by class; shading denotes 95% confidence intervals. **(c)** Spatial null distributions of the cross-species correlations in **b** (top: Mouse MERFISH 1, *P_null_* = 0.001; bottom: Mouse MERFISH 2, *P_null_* = 0.002, *n* = 1,000 rotations). **(d)** Cell-type marker-gene enrichment of AV-pole (*n* = 442) and PD-pole (*n* = 428) genes across seven cortical cell-type classes. AV-pole genes were significantly enriched for OPC, Astro, Micro, Neuro-In, and Neuro-Ex markers; PD-pole genes were significantly enriched for Oligo, Neuro-Ex, and Neuro-In markers. Dashed line, *P_FDR_* = 0.05; bubble size, enrichment score. \**P* < 0.05; \*\**P* < 0.01; \*\*\**P* < 0.001. Ex, excitatory; In, inhibitory; Glia, glial; Vasc, vascular; Oligo, oligodendrocyte; OPC, oligodendrocyte progenitor; Astro, astrocyte; Micro, microglia; Per, pericyte; Neuro-Ex, excitatory neuron; Neuro-In, inhibitory neuron.

**Extended Data Figure 3.**
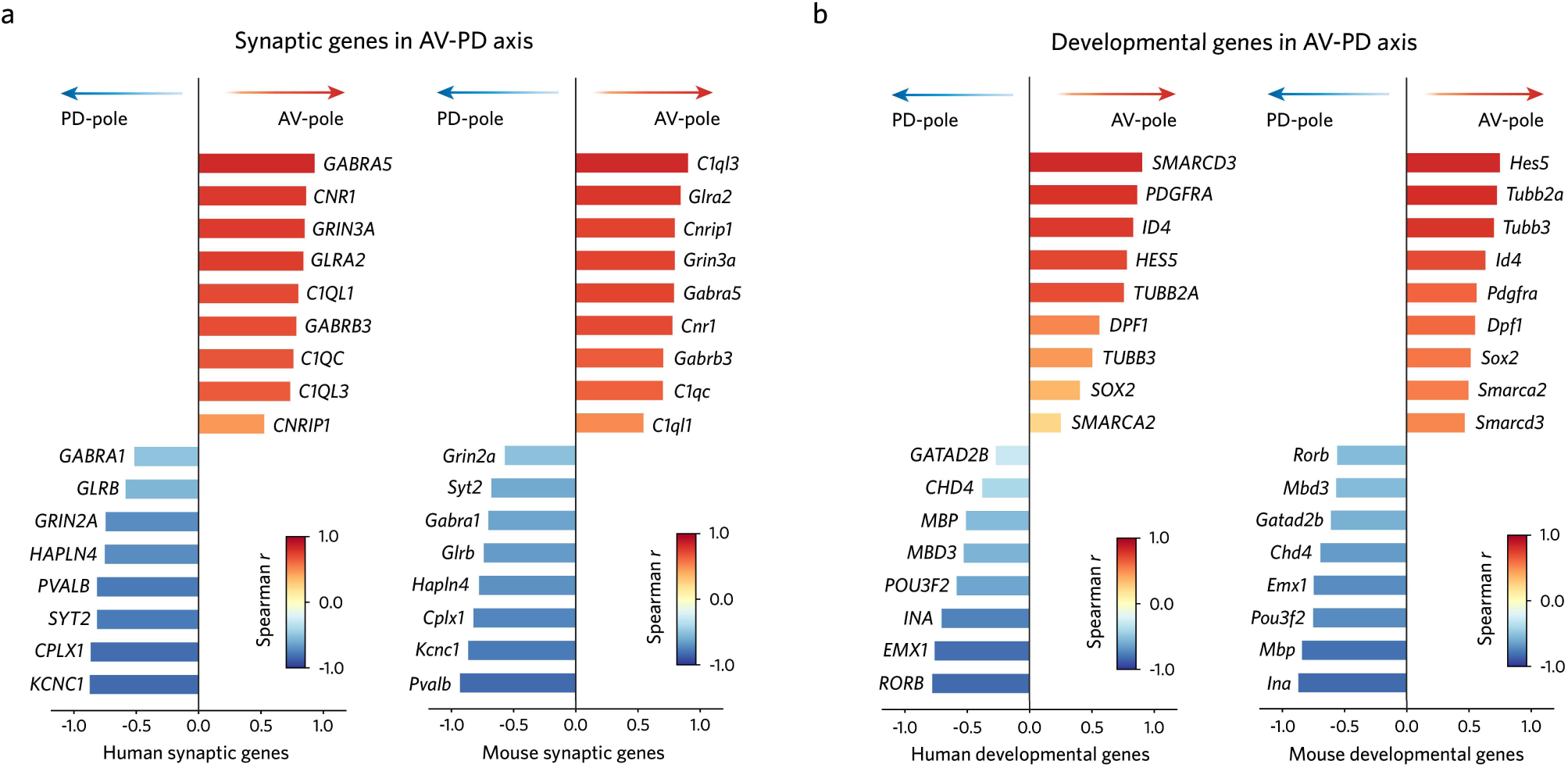
Canonical synaptic and developmental genes polarize along the AV–PD axis in humans and mice. **(a)** Spearman correlations between the AV–PD axis and cortical expression of representative synaptic genes in humans (left) and mice (right). Positive and negative correlations indicate AV- and PD-pole bias, respectively; colors indicate Spearman *r*. **(b)** As in a, for representative developmental genes.

**Extended Data Figure 4.**
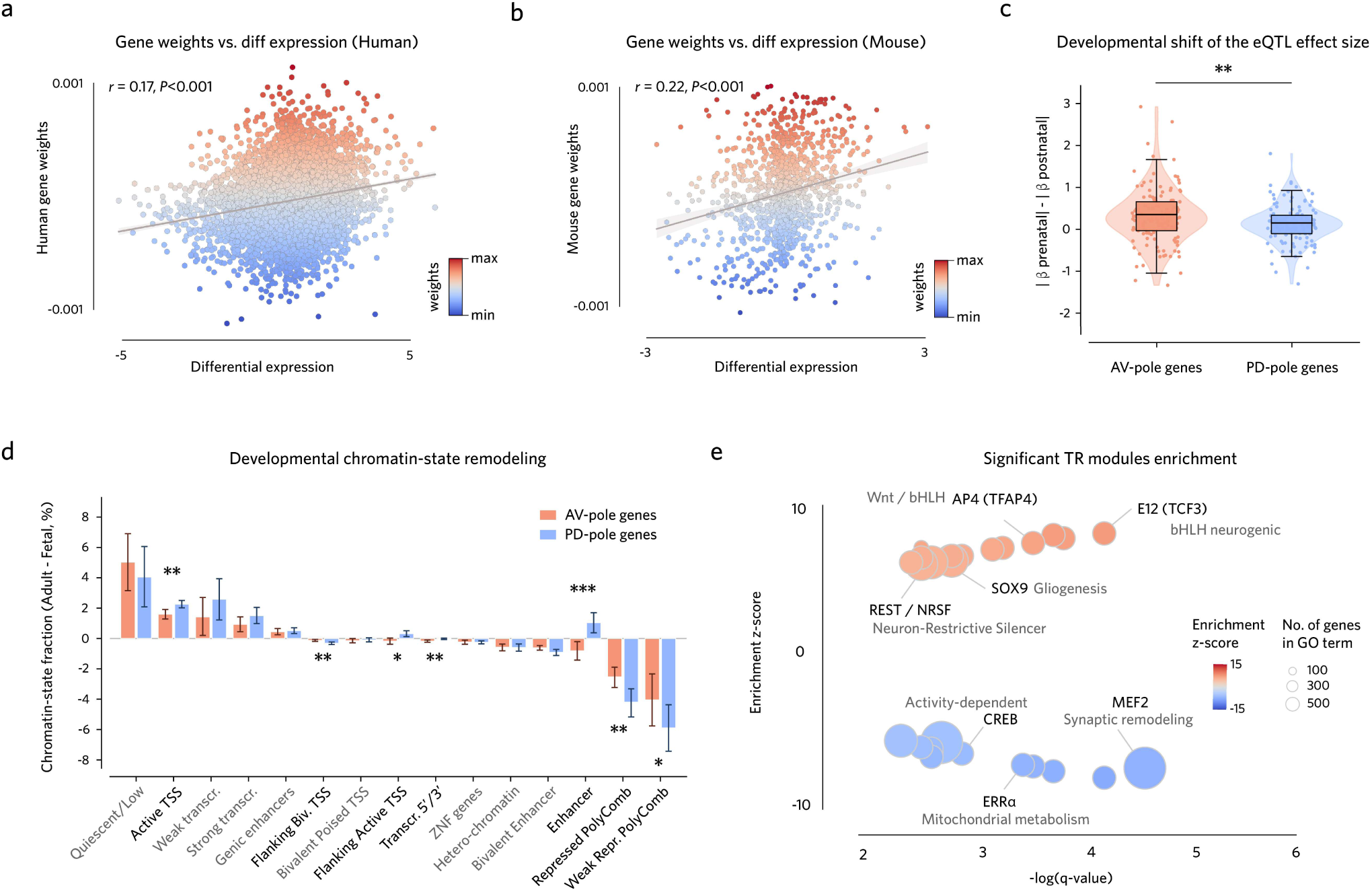
Regulatory features associated with AV–PD developmental timing. **(a)** AV–PD gene weights versus prenatal to postnatal differential expression in humans (Pearson *r* = 0.17, *P* = 1.69 × 10^−65^, *n* = 11,622 genes). Points are colored by gene weight; shading denotes the 95% confidence interval of the linear fit. **(b)** As in **a**, for mice (*r* = 0.22, *P* = 9.85 × 10^− 15^, *n* = 1,157 genes). **(c)** Developmental shift of *cis*-eQTL effect sizes (|β_prenatal| − |β_postnatal|) for AV-pole (*n* = 129) and PD-pole (*n* = 127) genes. AV-pole genes showed greater prenatal-biased *cis*-eQTL effects (rank-biserial *r* = 0.18, *P* = 0.006; two-sided Mann–Whitney *U* test). Violin plots show kernel density; box, median and 25th–75th percentiles; whiskers, 1.5 × IQR. **(d)** Developmental changes in chromatin-state fractions (Adult − Fetal, %) for AV-pole (*n* = 442) and PD-pole (*n* = 428) genes. Bars show mean ± s.e.m.; differences were assessed using two-sided Mann–Whitney *U* tests with FDR correction. **(e)** Transcriptional-regulator (TR) module enrichment for AV-pole (*n* = 442) and PD-pole (*n* = 428) genes. Positive and negative enrichment z-scores indicate AV- and PD-pole enrichment, respectively; bubble size, number of genes per TR module. \**P* < 0.05; \*\**P* < 0.01; \*\*\**P* < 0.001. eQTL, expression quantitative trait locus; TR, transcriptional regulator.

## Methods

### Human and mouse cortical atlas

The human cortical parcellation was based on a Brainnetome atlas derived from structural connectivity^78^. The left-hemisphere atlas was mirrored to the right hemisphere to ensure symmetry across the whole cortex. The final human cortical atlas consists of 105 cortical ROIs in standard MNI152 space. The mouse cortical atlas was obtained from the Allen Mouse Brain Atlas^79^. Small brain regions were aggregated into larger regions using hierarchical clustering to reduce fragmentation. The final mouse atlas is in CCFv3 space and comprises 39 cortical regions.

### Human transcriptomic data preprocessing

#### AHBA dataset

Human gene expression data were obtained from the Allen Human Brain Atlas^13^ (AHBA). The data were downloaded via the Allen Institute API (https://human.brain-map.org) and preprocessed using the abagen package^80^. During preprocessing, we used data from six donors, each containing log2 expression values of 58,692 gene probes derived from multiple tissue samples. For probe selection, we applied differential stability (diff_stability). To retain as many genes as possible, no intensity-based filtering thresholds were applied. In addition, to increase spatial coverage of the samples, the lr_mirror parameter was set to bidirectional. For each donor, we applied scaled robust sigmoid normalization to standardize gene expression both within and across samples. The norm_matched parameter was set to False, and region_agg was set to None. All other parameters were kept at their default values. The final gene expression matrices were obtained using the get_expression_data function in the abagen package. After assignment, the gene expression matrices, which included samples from different donors, were z-score normalized at the sample level and then averaged across all samples to obtain the final gene expression vectors representing each brain region for spatial transcriptional gradient analysis.

#### BrainSpan dataset

The BrainSpan dataset^1,52^, after batch-effect correction and normalization, was downloaded from the Allen Institute website (https://www.brainspan.org). Cortical brain regions were retained, resulting in developmental data covering 15 cortical regions from 8 post-conception weeks (pcw) to 40 years of age. For spatial gradient analysis, gene expression matrices were z-score normalized at the brain-region level within each stage. For temporal expression analysis, spatially unnormalized matrices were concatenated across developmental stages and then z-score normalized across stages to capture each gene’s relative developmental expression profile.

### Mouse transcriptomic data preprocessing

#### ISH dataset

The energy volumetric gene expression data^27^ (including coronal and sagittal experiments) were obtained from the Allen Institute API (https://mouse.brain-map.org) as sequences of 32-bit floating-point values. These arrays were reconstructed into 3D medical images in NetCDF (MINC) format, using the RAS coordinate system (Right–Anterior–Superior) with the origin defined at the midline intersection of the anterior commissure. Coronal and sagittal datasets were processed independently in Python. To harmonize the two datasets, only sagittal genes that were also present in the coronal dataset were retained. The images were imported using the nilearn library, masked, reshaped, and converted into a voxel × gene matrix. Expression values were log2 transformed. For genes with multiple hybridization experiments, voxel-wise averages were calculated. Genes with missing values in more than 20% of voxels were removed, and the remaining missing values were imputed using a K-nearest-neighbors (KNN) approach. The Allen CCFv3 atlas image was registered to the masked volumetric images, assigning each voxel an independent brain region label. Cortical brain regions were retained, and gene expression was z-score normalized at the voxel level and then averaged, resulting in a brain region × gene expression matrix for spatial transcriptional gradient analysis.

#### Spot array dataset

The whole-brain spot array sequencing data^28^ from three adult male mice were preprocessed, including log2 transformation and batch-effect correction across animals, and subsequently aligned to the CCFv3 standard space. In the processed gene expression matrix, each row corresponds to a spot, which is indexed by a unique label from the Allen CCFv3 hierarchical atlas. Spatial annotations were updated according to the hierarchical relationships between brain regions in the mouse brain atlas we used. After z-score normalization at the spot level, the data were averaged to obtain a brain region × gene expression matrix for spatial transcriptional gradient analysis.

#### MERFISH dataset

The quality-controlled and normalized mouse MERFISH spatial transcriptomic gene expression data^29,30^ were downloaded from the Allen Institute API (https://portal.brain-map.org/atlases-and-data/bkp/abc-atlas). The segmented cell coordinate index labels were reassigned according to the hierarchical structure of the mouse brain atlas used in our study. After z-score normalization at the segmented cell level, the data were averaged to obtain a brain region × gene expression matrix for spatial transcriptional gradient analysis.

#### Developing mouse brain atlas

The in situ hybridization (ISH) data from the Allen Developing Brain Atlas^26^ are organized in a grid-based format. Gene expression within each grid voxel is quantified as expression energy, defined as the product of the mean signal intensity and the fraction of expressing pixels. The spatial resolution ranges from 80 to 200 μm, depending on the developmental stage. We used datasets from seven developmental time points: E13.5, E15.5, E18.5, P4, P14, P28, and P56. Each dataset additionally provides a corresponding reference grayscale template and anatomical annotation labels. Grid data corresponding to the pallium were retained for analysis. Within each developmental stage, genes with zero expression across all grid voxels were removed, and the intersection of genes across all stages was used for downstream analyses. Expression values were log₂-transformed. For spatial gradient analysis, gene expression matrices were z-score normalized at the grid level within each stage. For temporal expression analysis, spatially unnormalized matrices were concatenated across developmental stages and then z-score normalized across stages at the grid level to capture each gene’s relative developmental expression profile.

### Cell-type spatial distribution dataset

#### Macaque-derived simulated human atlas

A whole-cortex human cell-type atlas, simulated from macaque spatial transcriptomics^31^, was obtained from Dong et al. (https://github.com/Dthbca/HomoloMap). The cross-species mapping pipeline comprised three key steps. First, homologous cell types were identified between human and macaque using MetaNeighbor, a framework that quantifies cross-species cell-type replicability based on the similarity of transcriptional signatures^81^. Second, a connectome-based joint embedding framework^50^ was applied to construct a shared mapping space across species, enabling projection of macaque cell-type composition spatial distributions onto the human cortex. Third, for each cortical region in the Brainnetome atlas^78^, the relative proportion of each cell type was computed with respect to all cell types, followed by z-score normalization across the whole cortex, yielding spatial ratio maps for 23 cell types.

#### Human single-nucleus dataset

The large-scale single-nucleus transcriptomic dataset of the human brain was provided by the Linnarsson laboratory^32^. The dataset comprises cell-type counts from three adult human donors, spanning approximately 32 anatomically defined regions across the cortex. We restricted the analysis to cortical regions and retained only the 23 cell types at the supercluster level used in this study. Similarly, for each specific cell type, its relative proportion in each cortical region was computed with respect to all cell types, followed by z-score normalization across the cortex.

#### Mouse MERFISH atlas

Mouse cell metadata with parcellation annotations were obtained from the Allen Institute API (https://portal.brain-map.org/atlases-and-data/bkp/abc-atlas), as reported by Zhang et al.^29^ and Yao et al.^30^. The metadata include cell-type annotations and CCFv3 region labels derived from MERFISH image segmentation coordinates. Subcluster-level cell labels were aggregated into 23 cell types based on the hierarchical organization of cell classes used in this study. For each specific cell type, its relative proportion in each cortical region was computed with respect to all cell types, followed by z-score normalization across the cortex.

#### Transcriptomic decomposition and gene weight characterization

The human cortical transcriptional gradient was published by Huang et al.^24^. This gradient is derived from the first principal component of region-specific transcriptional embeddings, explaining 26.8% of the variance. The mouse cortical transcriptional gradient was independently decomposed from four different transcriptomic datasets^27–30^ using PCA. For downstream analyses, we selected the gradient map derived from the Spot array dataset^28^. Although in this dataset the target gradient corresponds to the second principal component, explaining 13.2% of cortical variance, it includes a rich set of over 10,000 candidate genes for subsequent analyses. Mouse genes were mapped to their human homologues using the NCBI HomoloGene database^82^, and then intersected with genes from the AHBA dataset to construct a shared gene set for cross-species analyses, yielding a totally 12,731 genes.

To identify the weights of genes contributing to the transcriptional gradient, we employed a LASSO-PCR pipeline. This method preserves co-expression networks while enabling sparse feature selection^83^. Specifically, the gene expression matrix was subjected to principal component analysis (PCA) to reduce dimensionality and capture major axes of variation across cortical regions. LASSO regression was then applied in the PCA space to predict the transcriptional gradient. To obtain gene-level weights in the original gene space, the regression coefficients from the LASSO model were projected back through the PCA components as follows:

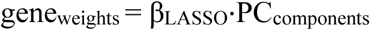

where β_LASSO_ represents the LASSO coefficients in PCA space and PC_components_ denotes the PCA loading matrix. The above LASSO-PCR pipeline was applied independently to human AHBA expression data^13^ and mouse Spot-array expression data^28^ to obtain gene weights in each species. These gene weights were subsequently used for downstream analyses, including developmental spatial pattern mapping.

#### Statistical testing of spatial correlation

In humans, we assessed the statistical significance of whole-brain comparisons while accounting for spatial autocorrelation, using the *vazquez_rodriguez* method, an Alexander-Bloch adaptation for parcellated data implemented in the neuromaps toolbox^84^. This method projects parcel centroids onto the cortical surface and reassigns them by minimizing Euclidean distance, thereby generating spatially constrained null distributions. In mice, the transcriptional axis and the maps used for comparisons were projected onto human space using the TransBrain framework^24^, and the significance of spatial autocorrelation was assessed using the same procedure as for human maps.

#### Enrichment analysis of cross-species axis-related genes

Enrichment analyses were performed using a hypergeometric test. Specifically, genes significantly associated with the transcriptional axis in both human and mouse were first identified based on Spearman correlation and spatial spin tests (FDR-corrected *P* < 0.05). These genes were further stratified according to the sign of the correlation coefficient into AV-biased genes (*r* > 0; preferentially expressed in anterior– ventral regions; human: 2,784 genes; mouse: 1,376 genes) and PD-biased genes (*r* < 0; preferentially expressed in posterior–dorsal regions; human: 2,123 genes; mouse: 1,808 genes). Genes showing consistent significant associations across species were further selected (AV-pole genes: n = 442; PD-pole genes: n = 428) as target genes (**Supplementary Table 4**). Hypergeometric enrichment analysis was then performed separately for AV-pole and PD-pole target gene sets across functional modules. Enrichment was assessed using a one-sided hypergeometric test with the 12,731 AHBA-detected homologous genes as the background population. For each term, the marker gene set was restricted to genes detected in the 12,731 homologous genes. The upper-tail probability was calculated as scipy.stats.hypergeom.sf (k−1, N, K, n), where N = 12,731, K is the number of AHBA-detected genes annotated to the term, n is the size of the target gene list, and k is the observed overlap. The enrichment score was defined as the observed-to-expected ratio (k/E, where E = K·n/N). P-values were corrected for multiple comparisons using the Benjamini–Hochberg false discovery rate (FDR). This approach enables the investigation of biological processes associated with gradient-related genes and the identification of module-specific “hit” genes, thereby facilitating functional interpretation and downstream analyses, such as gene expression trajectory profiling within selected gene subsets.

### Linking axis-related genes to biological processes and synaptic functions

#### Gene ontology enrichment

We performed pathway and Gene Ontology (GO) enrichment analyses using the Metascape platform^85^ (https://metascape.org/) to identify biological processes overrepresented among genes associated with the transcriptional axis. Significantly enriched terms after multiple-comparison correction are reported in **Supplementary Table 5**.

#### Synaptic function enrichment

We performed synaptic function enrichment analysis against the SynGO v1.3 ontology, a manually curated, expert-reviewed annotation of synaptic biology^86^. To avoid redundancy across the hierarchy, we restricted the analysis to five top-level biological-process terms — presynaptic process (SYNGO:presynprocess), postsynaptic process (SYNGO:postsynprocess), synapse organization (GO:0050808), synaptic metabolism (SYNGO:metabolism), and synaptic transport (SYNGO:transport). Enrichment of the AV-pole and PD-pole gradient-related gene sets within each term was assessed using the hypergeometric test described above, with the 12,731 AHBA-detected homologous genes as the background, and *P* values were corrected for multiple comparisons using the Benjamini–Hochberg false discovery rate (FDR).

### Linking cross-species transcriptional axis to cortical cell types

#### Cell-type spatial distribution alignment

The spatial pattern of each of the 23 cell-type proportions in human and mouse was independently correlated (Spearman *r*) with the transcriptional axis, yielding two vectors that quantify the associations between the transcriptional axis and cell-type spatial distributions in each species, referred to as cell-type– gradient association profiles. In parallel, within each species, we computed the pairwise spatial autocorrelation matrix (Spearman *r*) across the 23 cell types. A diffusion-map embedding algorithm, implemented in the brainspace toolbox^51^ (α = 0.8, sparsity = 0.5), was then applied to these matrices to derive low-dimensional embedding vectors, capturing the intrinsic organization of cell-type spatial distributions within each species, referred to as intrinsic cell-type spatial organization profiles. Pearson correlation was used to quantify the similarity between these profiles. In addition, we also assessed whether the 23 cell types could support cross-species prediction of the transcriptional axis. Specifically, partial least squares (PLS) regression was trained on human data to model the relationship between cell-type spatial distributions and the transcriptional axis. The learned weights were then transferred to the mouse cell-type spatial maps to predict the mouse transcriptional axis.

To determine whether the cross-species similarity of cell-type–gradient association profiles was specifically driven by the transcriptional axis rather than generic spatial similarity, we performed permutation testing. In each iteration, the transcriptional axis was spatially rotated using the *vazquez_rodriguez* method—an Alexander-Bloch adaptation for parcellated data implemented in the neuromaps toolbox^84^—to generate null maps. These maps were subsequently projected onto the mouse cortex using the TransBrain framework^24^, yielding spatially homologous but transcriptionally unrelated gradients across species. Cell-type–gradient association profiles were then recomputed based on these null gradients, and their cross-species similarity was reassessed to construct a null distribution. This procedure was repeated 1,000 times.

#### Cell-type marker-gene enrichment

Cell type-specific gene sets were sourced from Seidlitz et al.^33^. Enrichment of the AV-pole and PD-pole gradient-related gene sets within each term was assessed using the hypergeometric test described above, with the 12,731 AHBA-detected homologous genes as the background, and *P* values were corrected for multiple comparisons using the Benjamini–Hochberg false discovery rate (FDR).

#### Linking cross-species transcriptional axis to macroscale phenotypes

The principal-component gradients of thalamocortical projections in humans and mice were obtained from Oldham et al.^34^. The human T1/T2-weighted (T1w/T2w) gradient was downloaded from the neuromaps toolbox^84^, originally published by Burt et al.^15^, whereas the mouse T1/T2w gradient was derived from Fulcher et al.^35^. The human excitatory/inhibitory (E/I) gradient was obtained from Hansen et al.^36^, and the proxy for the mouse E/I ratio was estimated from the relative proportions of excitatory and inhibitory neurons in cortical regions from the MERFISH dataset^29,30^. All voxel-wise or atlas-based human macro-scale phenotypes were aligned to MNI 152 space. The mean value of voxels within each region defined by the Brainnetome atlas^78^ was extracted. Associations between the transcriptional axis and macroscale phenotype measures were quantified using Spearman correlation.

### Spatial development topology of the AV–PD axis

#### Human developmental gradient patterns

To examine how the adult gradient evolves over development, gene expression data at each stage of the BrainSpan dataset^1,52^ were weighted by gene weights derived from the adult transcriptional gradient to reconstruct spatial patterns.

#### Mouse postnatal gradient patterns

As in humans, gene expression data at each mouse postnatal stage were weighted by gene weights derived from the adult transcriptional gradient. To enable visualization and region-level quantitative comparisons, the low-resolution grid data were registered to high-resolution grayscale reference templates (antsRegistration, ANTs). We further utilized the DevCCF atlas^87^ to obtain fine-grained cortical annotations. This atlas provides high-resolution light-sheet fluorescence microscopy templates and corresponding CCFv3 anatomical labels for postnatal stages P4, P14, and P56. For each developmental stage, the high-resolution light-sheet templates were registered to the corresponding grayscale reference templates used for visualization, and the annotated CCFv3 label images^79^ were transformed accordingly (antsRegistration, ANTs). Due to the absence of annotations for P28, the high-resolution light-sheet template from P56 was registered to the P28 grayscale reference template. After registration, voxel-wise gradient values were aggregated to cortical regions based on the mouse brain atlas used in this study, enabling quantitative comparisons of cortical gradient similarity across developmental stages.

#### Mouse prenatal gradient patterns

Similarly, gene expression data at each prenatal stage were weighted by gene weights derived from the adult transcriptional gradient. The resulting prenatal gradient maps were registered to high-resolution grayscale reference templates for visualization. However, neither the Allen Developing Brain Atlas^26^ nor the DevCCF atlas^87^ provides anatomical annotations aligned to the CCFv3 framework for prenatal stages. Notably, we observed that postnatal and adult gradient patterns consistently exhibit a spatial organization along an anterior–ventral to posterior–dorsal axis on the midsagittal plane, corresponding to the intrinsic geometric organization of the brain. To enable quantitative assessment of prenatal transcriptional gradients, we therefore computed a geometric gradient based on the three-dimensional coordinates of the midsagittal plane using the brainspace toolbox^51^. This geometric gradient was used to partition the brain into anterior– ventral and posterior–dorsal domains. Based on this partition, we assessed whether prenatal stages exhibit gradient patterns consistent with those observed postnatally, and identified genes that maintain differential expression between anterior–ventral and posterior–dorsal regions throughout development.

### Developmental timing analysis of the AV–PD axis

#### Regional developmental slope matrix decomposition

For both human and mouse developmental datasets^1,26,52^, gene expression matrices without spatial normalization were first concatenated across developmental stages, followed by z-score normalization across the whole cortex across stages to capture the relative expression strength of each gene over development. To control for global expression biases introduced by differences in regional volume across mouse developmental stages, the mean expression across all genes within each grid was regressed out from the z-score–normalized mouse gene expression matrix. No such regression was applied to the human gene expression data. For the BrainSpan dataset^1,52^, due to the relatively dense sampling of time points, the entire developmental period was divided into seven stages: 8–13 pcw, 16–19 pcw, 21–26 pcw, 4 months–2 years, 3–8 years, 13–15 years, and 18–40 years. Subsequently, gene expression data were aggregated by developmental stages and brain regions through averaging. For each cortical region and each gene, the normalized expression change from the earliest detectable prenatal stage to adulthood was computed and defined as the developmental slope:

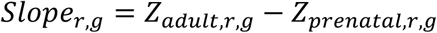

where ***r*** represents the cortical region and ***g*** represents the gene. This generated a regional × gene developmental slope matrix, which captures genome-wide patterns of developmental transcriptional changes across cortical regions. PCA decomposition was applied to the human and mouse matrix. The first principal component (PC1) was defined as the developmental slope gradient.

#### Gene expression trajectories

For a given gene set *G*, the developmental expression trajectory at stage t is defined as:

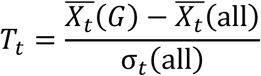

where: 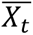(*G*) is the mean expression of the selected gene set G at stage t; 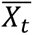(all) is the mean expression of all genes at stage t; σ_t_(all) is the standard deviation of expression of all genes at stage t. To visualize the developmental expression trajectories of axis-related gene sets, the following procedure was applied separately for AV (Spearman *r* > 0) and PD-biased (Spearman *r* < 0) gene sets in both human and mouse datasets. At each iteration, 200 genes were randomly sampled with replacement from the corresponding gene set, and their stage-wise trajectory was computed as defined above. To reduce stage-to-stage noise and emphasize developmental trends, the resulting trajectory was smoothed using locally weighted scatterplot smoothing (statsmodels.nonparametric.smoothers_lowess; smoothing parameter frac = 0.6) over the seven developmental stages, and further interpolated to 200 evenly spaced points along the developmental axis using a cubic B-spline (scipy.interpolate.make_interp_spline). This sampling– smoothing–interpolation procedure was repeated 1,000 times to obtain a distribution of trajectories. To assess whether AV- and PD-biased gene sets exhibited significantly distinct developmental trajectories, a trajectory-level label-permutation test was performed separately in human and mouse datasets. At each permutation, gene identities were randomly reassigned between AV- and PD-biased gene sets while preserving the original group sizes. The same sampling–smoothing–interpolation procedure described above was then repeated to generate group-level developmental trajectories for each permuted gene set. Trajectory differences were quantified as the summed squared distance between the mean trajectories across all interpolated developmental points:

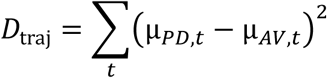

where (*y̅*_t_) denotes the mean expression trajectory at developmental position (t). This procedure was repeated 1,000 times to generate a null distribution of trajectory differences. Empirical significance was determined as the proportion of permutations yielding a trajectory distance greater than or equal to the observed value. To further compare the overall developmental expression trends of AV- and PD-biased gene sets, we calculated the cumulative distributions of developmental expression changes for the two gene sets in both humans and mice. Within each gene set, genes were ranked by their developmental slope, and the cumulative fraction of genes below a specific slope value was calculated to generate empirical cumulative distribution curves. A leftward shift of the curve indicates enrichment of genes with negative developmental changes, reflecting a tendency for expression decline toward adulthood.

#### Binned characterization of developmental gradient in AV–PD spatially biased genes

To assess whether genes distributed along the AV–PD axis exhibit graded developmental expression slopes, we performed a binned analysis of developmental gradients based on spatial bias independently in humans and mice. In humans, gene-wise cortical spatial bias scores were calculated as the Spearman correlation between AHBA^13^ adult cortical expression patterns and the AV–PD axis. In mice, the same approach was applied using spot-array spatial transcriptomic data^28^. Genes were then ranked from AV-biased to PD-biased according to their spatial bias scores and divided into 10 continuous bins within each species. Importantly, gene binning was defined solely by adult gene spatial expression patterns and was independent of developmental expression. Developmental slopes were subsequently calculated for each gene within each bin as described above, and the mean developmental slope of each bin was used to characterize the developmental slope gradient along the AV–PD axis.

#### Developmental eQTL analysis

To test whether AV-pole and PD-pole genes differ in the prenatal–postnatal shift of cis-eQTL effects, we used the BrainVar dorsolateral prefrontal cortex eQTL resource (Werling et al.^88^, Table S5, HCP_allEQTLs_FDR0.05 sheet), which provides all significant SNP−cis-eQTL pairs (*FDR* < 0.05) together with their prenatal- and postnatal-specific effect sizes (Beta_Pre, Beta_Post). For each SNP, we computed the prenatal–postnatal shift in cis-regulatory effect magnitude as Δ = |Beta_Pre| − |Beta_Post|, such that Δ>0 indicates stronger cis-eQTL during prenatal development. Because individual genes are typically associated with multiple cis-eQTLs, we aggregated to a single per-gene value by taking the median Δ across all cis-eQTLs assigned to that gene. Then, the Δ distributions were compared between AV-pole and PD-pole genes using a two-sided Mann–Whitney *U* test. Effect size was reported as the rank-biserial correlation *r* = 2U / (n₁n₂) − 1, with positive r indicating a larger prenatal–postnatal shift in AV-pole genes.

#### Developmental chromatin state analysis

The 15-state chromatin state segmentation files used in this study were downloaded from the Roadmap Epigenomics data portal (https://egg2.wustl.edu/roadmap/data/byFileType/chromhmmSegmentations/ChmmModels/coreMarks/jointModel/final). These 15 chromatin states were originally inferred by ChromHMM^89,90^, which applies an unsupervised hidden Markov model to five histone modification ChIP-seq tracks (H3K4me3, H3K4me1, H3K27me3, H3K36me3, H3K9me3), yielding per-interval chromatin state annotations across the genome. Fetal brain was represented by E081 (male fetal brain) and E082 (female fetal brain), and adult brain by E067 (angular gyrus), E068 (anterior caudate), E069 (cingulate gyrus), and E073 (dorsolateral prefrontal cortex). All segmentation files are provided in the hg19/GRCh37 human genome assembly.

The chromatin state composition of each AV-pole and PD-pole gene was quantified as the proportion of base pairs within the gene’s TSS-centered window that were assigned to each of the 15 chromatin states. Specifically, for each gene, the promoter region was defined as a 10 kb window centered on the transcription start site (TSS; ±5 kb), based on GENCODE v19 annotation^91^. TSS coordinates were assigned in a strand-aware manner: the gene start coordinate was used for plus-strand genes, and the gene end coordinate for minus-strand genes. For genes with multiple annotated transcripts, the canonical TSS of the principal transcript was used. Within this window, ChromHMM segments were intersected with the window, and the fraction of base pairs assigned to each of the 15 chromatin states was computed, yielding for each gene a 15-dimensional fractional coverage vector summing to 1.

Per-gene composition vectors were then averaged across the two fetal samples (E081, E082) and across the four adult samples (E067, E068, E069, E073) to obtain stage-specific composition vectors. For each gene and each of the 15 chromatin states, a developmental shift was computed as Δ = adult − fetal, with positive Δ indicating that the state is more prevalent in adult than fetal brain. To compare developmental chromatin-state differences between the AV-pole and PD-pole genes, the per-gene Δ values were contrasted between AV-pole and PD-pole genes using a two-sided Mann–Whitney *U* test, with the rank-biserial correlation *r* = 2U / (n₁n₂) - 1 reported as the effect size. P-values were corrected for multiple comparisons across the 15 states using the Benjamini–Hochberg false discovery rate (FDR), and states with *FDR* < 0.05 were considered to show significantly different developmental shifts between the two gene groups.

#### Transcription factor enrichment

We performed transcription factor (TF) motif enrichment analyses using the Metascape platform^85^ (https://metascape.org/) to identify upstream regulators overrepresented among genes associated with the transcriptional axis. AV-pole and PD-pole gene sets were submitted separately, and enrichment was tested against the integrated TRRUST and TF-target motif libraries using the default hypergeometric framework with Benjamini–Hochberg FDR correction.

### Linking the AV–PD axis to autism risk

#### Autism transcriptional development deviations

The autism cortical developmental transcriptomic dataset was obtained from Schwarz et al.^37^, who performed single-nucleus RNA sequencing of the whole cortex across developmental stages (E14.5, P4, and P14) in 11 autism-associated mouse genotypes and wild-type controls, followed by standard preprocessing. Because this dataset lacks fine-grained cortical regional annotations, we used previously defined AV–PD spatial bias gene bins as a reference framework. Expression values were first z-score normalized across all developmental stages and genotypes, including wild-type controls, to place trajectories on a common scale. Autism models were initially pooled into a single ASD group, and gene-wise developmental slopes were calculated for the WT and ASD groups as described above. The resulting slope distributions were then compared within each AV–PD spatial bias bin. The same analysis was subsequently repeated for individual autism genotypes. Finally, we quantified ASD-associated developmental slope differences relative to WT within each bin and assessed their spatial deviations along the AV–PD axis.

#### Autism brain volume deviations

Mouse structural MRI data of autism mouse models were obtained from Ellegood et al.^38^. All animals were transferred to the Mouse Imaging Centre (Toronto, Canada) for harmonized ex vivo neuroimaging. After excluding samples with missing genotype information and mutations affecting the Y chromosome, the final dataset comprised cohorts spanning 13 distinct ASD risk genes. Regional brain volumes were quantified from the structural MRI data using deformation-based morphometry. In brief, all images were first corrected for intensity non-uniformities and subsequently aligned to the unbiased global average template provided by the dataset. Registration was performed using ANTs^92^, with cascaded linear (6-parameter rigid followed by 12-parameter affine) and nonlinear (SyN diffeomorphic) stages. From the resulting transformations, Jacobian determinant maps encoding the local volume change of each voxel relative to the template were computed in template space. Our mouse cortical atlas was registered to the same template, and per-region volumes were derived by summing Jacobian values within each cortical ROI. Finally, ROI volumes were normalized by the mean ROI volume across the cortex to remove global volume effects.

For the computation of mutation-specific brain volume deviation maps, following Ellegood et al.^38^ and given the limited sample size per genotype, age and sex were not regressed out. For each genotype, mutation-specific brain volume deviation maps were computed as Cohen’s d effect sizes between mutant and wild-type mice for each cortical ROI. Next, to identify subgroups of ASD mutations that share similar patterns of cortical volume alteration, we computed the pairwise Spearman rank correlation matrix across the 13 genotype-specific deviation maps and applied hierarchical clustering to this similarity matrix. Within each resulting cluster, deviation maps from all member genotypes were averaged and the mean map was z-score normalized across cortical regions, yielding a cluster-level cortical volume alteration profile. The spatial correspondence between each cluster-level profile and the cross-species transcriptional axis was then quantified using Spearman correlation, with statistical significance assessed against spatial autocorrelation-preserving null distributions as described above.

#### Psychiatric risk-gene enrichment

Psychiatric risk associations of AV-pole and PD-pole genes were assessed using three complementary lines of evidence: (1) common-variant risk from genome-wide association studies (GWAS), via H-MAGMA; (2) differentially expressed genes (DEGs) from postmortem brain transcriptomics; and (3) curated autism risk genes from the SFARI Gene database.

**(1) H-MAGMA:** we performed common-variant gene-set association analysis using H-MAGMA^93^, which extends MAGMA^94^ by leveraging chromatin-conformation (Hi-C) contacts to assign non-coding SNPs to their physical target genes, rather than relying solely on linear genomic proximity. SNP-level summary statistics were obtained from the Psychiatric Genomics Consortium (PGC) GWAS of autism spectrum disorder^39^ and schizophrenia^40^. Gene-level statistics were computed using the 1000 Genomes European reference panel^36^ (g1000_eur) and two precomputed H-MAGMA SNP-to-gene annotations: the fetal-brain Hi-C annotation^95^ and the adult-brain Hi-C annotation^96^. AV-pole and PD-pole gene symbols were converted to Ensembl gene identifiers via GENCODE v19^91^ and formatted as MAGMA gene-set input files. For each disorder × developmental-stage combination (SCZ-fetal, SCZ-adult, ASD-fetal, ASD-adult), competitive gene-set analysis was performed, which tests whether the mean gene-level association statistic in the target gene set exceeds that of all other genes in the genome, accounting for gene size, gene density, and SNP-level linkage disequilibrium. Significance was reported as the one-sided competitive *P* value and the standardized regression coefficient β. *P*-values across the four disorders × stage combinations were corrected using the Benjamini–Hochberg false discovery rate (FDR).
**(2) Differential expression risk gene enrichment:** Disease risk genes were identified through case– control differential expression analysis of RNA-seq data from postmortem brain tissue, as follows: ASD (Parikshak et al.^41^, FDR-adjusted *P* value, ASD versus CTL < 0.05; Gandal et al.^44^, ASD.fdr < 0.05; Gandal et al.^46^, WholeCortex_ASD_FDR < 0.05); MDD (Jaffe et al.^45^, Cortex_adjPVal_MDD < 0.05); SCZ (Fromer et al.^42^, FDR estimate < 0.05; Jaffe et al.^43^, pval_adj < 0.05; Gandal et al.^44^, SCZ.fdr < 0.05); BD (Gandal et al.^44^, BD.fdr < 0.05). To improve robustness, for ASD and SCZ, only genes that were reproducibly identified as differentially expressed in at least two independent studies were retained as risk genes.
**(3) SFARI:** Rare-variant ASD risk genes were obtained from the SFARI Gene database^47^ (https://gene.sfari.org/), restricted to genes with SFARI gene scores of 1 (High Confidence) or 2 (Strong Candidate), thereby retaining evidence-supported risk genes.

DEG and SFARI enrichment of the AV-pole and PD-pole axis-related gene sets was assessed using the hypergeometric test described above, with the 12,731 AHBA-detected homologous genes as the background, and *P*-values were corrected for multiple comparisons using the Benjamini–Hochberg false discovery rate (FDR).

## Data availability

The Allen Human Brain Atlas (AHBA) gene expression data are available at https://human.brain-map.org. The BrainSpan developmental transcriptome can be accessed at https://www.brainspan.org. The Allen Mouse Brain ISH dataset and the developing mouse brain ISH atlas are available via the Allen Brain Atlas API at https://mouse.brain-map.org and https://developingmouse.brain-map.org, respectively. The whole-brain mouse spot-array spatial transcriptomic data can be accessed at https://www.molecularatlas.org. The two whole-brain mouse MERFISH datasets are available via the Allen Brain Cell Atlas at https://portal.brain-map.org/atlases-and-data/bkp/abc-atlas. The autism cortical development transcriptome can be accessed at https://adameykolab.hifo.meduniwien.ac.at/cellxgene/filecrawl/.2026_Nature_Schwarz. The single-nucleus dataset for the human brain is available at https://storage.cloud.google.com/linnarsson-lab-human. The Allen Mouse Brain Common Coordinate Framework (CCFv3) is available at https://atlas.brain-map.org, and the Developmental Common Coordinate Framework (DevCCF) atlas is available at https://kimlab.io/brain-map/DevCCF/. The Brainnetome atlas is available at https://atlas.brainnetome.org. The principal gradient of thalamocortical projections can be accessed from https://github.com/StuartJO/ThalamicGradients. The human T1w/T2w cortical myelin map is accessible through the neuromaps toolbox at https://github.com/netneurolab/neuromaps. The 15-state ChromHMM chromatin-state segmentations from the Roadmap Epigenomics Project are available at https://egg2.wustl.edu/roadmap/data/byFileType/chromhmmSegmentations. NCBI HomoloGene was used to map mouse–human homologous genes https://www.ncbi.nlm.nih.gov/homologene. The SynGO v1.3 synaptic gene ontology is available at https://www.syngoportal.org. The structural MRI dataset of autism mouse models is available at https://www.braincode.ca/content/public-data-releases#dr001. SFARI ASD rare-variant risk genes are available at https://gene.sfari.org. All other atlases and resources used in this study were obtained from previous publications, as detailed in the Methods section.

## Code availability

The pipeline for simulating whole-brain human cell-type spatial distributions from macaque spatial transcriptomic data is publicly available on GitHub at https://github.com/Dthbca/HomoloMap, under the MIT License. The data and pipeline used for the AV–PD axis analysis is publicly available at https://github.com/ibpshangzheng/cortical_axis, under the Apache License, Version 2.0 (Apache-2.0).

## Acknowledgements

This work was supported by the Science and Technology Innovation 2030-Brain Science and Brain-inspired Intelligence Project of China (2022ZD0211900) and Beijing Nova Program (Grant No.20250484761). The funders had no role in study design, data collection and analysis, decision to publish or preparation of the manuscript.

## Author contributions

A.L. and S.H. led the project. A.L., S.H. and Y.S. were responsible for the study conception and design. S.H. analyzed the data. S.H. and A.L. created the figures and wrote the paper. Y.S. aided in the writing of the paper. Y.P., X.T., C.D and T.Z. contributed to discussions of the results and the manuscript.

